# High spatial overlap but diverging age-related trajectories of cortical MRI markers aiming to represent intracortical myelin and microstructure

**DOI:** 10.1101/2022.01.27.477925

**Authors:** Olivier Parent, Emily Olafson, Aurélie Bussy, Stephanie Tullo, Nadia Blostein, Alyssa Salaciak, Saashi A. Bedford, Sarah Farzin, Marie-Lise Béland, Vanessa Valiquette, Christine L. Tardif, Gabriel A. Devenyi, M. Mallar Chakravarty

**Affiliations:** Computational Brain Anatomy (CoBrA) Laboratory, Cerebral Imaging Center, Douglas Mental Health University Institute, Montreal, Quebec, Canada; Integrated Program in Neuroscience, McGill University, Montreal, Quebec, Canada; Department of Psychiatry, McGill University, Montreal, Quebec, Canada; Department of Biomedical Engineering, McGill University, Montreal, Quebec, Canada; McConnell Brain Imaging Center, Montreal Neurological Institute, Montreal, Quebec, Canada; Department of Neurology and Neurosurgery, McGill University, Montreal, Quebec, Canada; Department of Radiology, Weill Cornell Medicine, New York, USA

**Keywords:** T1-weighted/T2-weighted ratio, Gray-white matter contrast, Boundary sharpness coefficient, Cortical thickness, Aging, Cortical Myelin

## Abstract

Cortical thickness (CT), gray-white matter contrast (GWC), boundary sharpness coefficient (BSC), and T1-weighted/T2-weighted ratio (T1w/T2w) are cortical metrics derived from standard T1- and T2-weighted magnetic resonance imaging (MRI) images that are often interpreted as representing or being influenced by intracortical myelin content. However, there is little empirical evidence to justify these interpretations nor have the homologies or differences between these measures been examined. We examined differences and similarities in group mean and age-related trends with the underlying hypothesis that different measures sensitive to similar changes in underlying myelo- and microstructural processes should be highly related. We further probe their sensitivity to cellular organization using the BigBrain, a high-resolution digitized volume stemming from a whole human brain histologically stained for cell bodies with the Merker stain.

The measures were generated on both the MRI-derived images of 127 healthy subjects, aged 18 to 81, and on the BigBrain volume using cortical surfaces that were generated with the CIVET 2.1.0 pipeline. Comparing MRI markers between themselves, our results revealed generally high overlap in spatial distribution (i.e., group mean), but mostly divergent age trajectories in the shape, direction, and spatial distribution of the linear age effect. Significant spatial relationships were found between the BSC and GWC and their BigBrain equivalent, as well as a correlation approaching significance between the BigBrain intensities and the T1w/T2w ratio in gray matter (GM) both sampled at half cortical depth.

We conclude that the microstructural properties at the source of spatial distributions of MRI cortical markers (e.g. GM myelin) can be different from microstructural changes that affect these markers in aging. While our findings highlight a discrepancy in the interpretation of the biological underpinnings of the cortical markers, they also highlight their potential complementarity, as they are largely independent in aging. Our BigBrain results indicate a general trend of GM T1w signal and myelin being spatially related to the density of cells, which is possibly more pronounced in superficial cortical layers.

**Highlights:**

– Different MRI cortical markers aim to represent myelin and microstructure
– These markers show high spatial overlap, but mostly divergent age trajectories
– It is unlikely that myelin changes are the source of the age effect for all markers
– Trend of MRI signal being related to cell density in more superficial cortical layers

## 1. Introduction

The microstructural organization of the human brain is defined by numerous anatomical organizations that include the cytoarchitecture (neuronal cell bodies), myeloarchitecture (organization of myelin sheaths), iron distribution, neuronal processes, vasculature, and glial cells (Bock et al., 2009; Eickhoff et al., 2005; Fukunaga et al., 2010; Tardif et al., 2016). T1-weighted (T1w) and T2-weighted (T2w) magnetic resonance imaging (MRI) contrasts represent a complex combination of these microstructural properties that is not yet completely understood (Tardif et al., 2016). The neurobiological specificity attributable to different intensities in T1w and T2w images is further confounded by experimental design choices such as MRI hardware and sequence acquisition parameters.

Histological studies have demonstrated that cortical T1w signal is influenced by both myelo- and cyto-architectural properties, although myelin was shown to be the main contributor to the contrast (Bock et al., 2009; Eickhoff et al., 2005). Other studies claimed that myelin was the largest contributor to quantitative T1 contrast, while iron was the largest contributor to quantitative T2* contrast (Stüber et al., 2014). However, iron and myelin largely colocalize in the cortex (Fukunaga et al., 2010), thus relating MRI signal mainly to myelo-architecture in both contrasts in the healthy cortex. In spite of these biophysical contrast mechanisms, several studies use metrics derived from the signal intensity as a representation of cortical “microstructure”, a non-specific term that does not have an agreed-upon biologically meaningful definition. Furthermore, metrics derived from T1w and T2w images are often interpreted as being influenced by myeloarchitecture, and more specifically the density or concentration of gray-matter (GM) myelin (Glasser & Van Essen, 2011; Olafson et al., 2021; Salat et al., 2009). In this manuscript, we seek to characterize the similarities and differences in three measures that are often used to describe cortical myelin and microstructure. These include: 1) The ratio of gray-to-white matter T1w signal intensities (gray-white matter ratio [GWC]; as originally proposed in Salat and colleagues (2009)), which is often used as a putative marker for myelination of deeper cortical layers (Chwa et al., 2020; Drakulich et al., 2021; Jørgensen et al., 2016; Vidal-Piñeiro et al., 2016); 2) The ratio of T1w to T2w signal (T1w/T2w ratio) has been proposed to be more sensitive to myelin than either contrast alone (Glasser & Van Essen, 2011; Grydeland et al., 2013, 2019), due to the generally inverse dependence of T1w and T2w signals on myelin; and 3) The boundary sharpness coefficient (BSC), recently proposed by our group, which examines the sharpness of the change in T1w signal intensity from superficial white matter (SWM) to gray matter (Olafson et al., 2021).

The BSC was inspired by methods used in cytoarchitectonic histological examinations in Avino & Hutsler (2010).

As previously mentioned, other neuroanatomical properties such as cortical iron and cell density have been shown to affect MRI signal (Eickhoff et al., 2005; Fukunaga et al., 2010), thus potentially also impacting the cortical measures previously mentioned.

Furthermore, SWM myelin could also have an impact, especially for the BSC and GWC markers, which sample intensities partly in the SWM. These other potential microstructural sources are rarely mentioned.

This myelin-specific dependency extends beyond cortical markers that directly measure the T1w signal. Morphological analyses relying on cortical thickness (CT) measures have also been shown to correlate with intracortical myelin (Natu et al., 2019; Patel et al., 2020; Shafee et al., 2015). Indeed, while CT was developed to assess cortical gray matter, it relies on the placement of the gray-white matter boundary on T1w images, which is typically established by algorithms as the location where the greatest change in contrast occurs (Salat et al., 2009), thus potentially depending on intracortical myelin density. Natu and colleagues (2019) found that CT reductions in the visual cortex observed during development were driven by myelination of deep cortical layers. Furthermore, studies reported a generally inverted correlation of CT and GM myelin across some areas of the cortex, particularly using “virtual histology” techniques relating cell-specific gene expression to MRI contrast (Patel et al., 2020; Shafee et al., 2015).

These cortical measures have also been reported as being sensitive to maturation and aging (Drakulich et al., 2021; Fjell et al., 2009; Grydeland et al., 2019; Olafson et al., 2021; Salat et al., 2009, 2011; Vidal-Piñeiro et al., 2016), suggesting that they are sensitive to normative age-related variations. However, for these measures to be meaningful in understanding neurobiology, and if they are indeed sensitive to the same myeloarchitectonic properties, their age-related trajectories should be similar. The most specific characterizations of intracortical myelin age trajectories have been done with quantitative R1 maps, an MRI contrast less biased by experimental choices and more specific to biophysical properties of the tissue (Marques et al., 2010; Tardif et al., 2016), and have found inverted U-shaped age trajectories across the cortex with earlier peaks in posterior regions (Erramuzpe et al., 2021). Therefore, we hypothesize that cortical markers representing intracortical myelin should follow similar age trajectories. Further, the influence of cytoarchitecture on these markers is not well characterized, even if cytoarchitecture is known to influence, to some degree, T1w signal (Eickhoff et al., 2005).

Thus, the overarching goal of this manuscript is to assess the similarities and differences characterized by these MRI-based cortical markers of morphology, microstructure, and myelin (i.e. CT, GWC, BSC, and T1w/T2w ratio). We first quantitatively compared the spatial distribution of these measures in a healthy population that spanned the adult lifespan. To assess if similar microstructural changes are at the source of age-related changes in all markers, we then compared their age-related trajectories between themselves. Additionally, we compare the spatial distribution of the markers with quantitative R1 maps. Next, using the BigBrain histological reconstruction (Amunts et al., 2013), we assessed whether these markers are also impacted by cyto-architectural organization by analyzing the homologies between cytoarchitectonic- and MRI-derived measures.

## 2. Methods

### 2.1 Participants

A total of 174 healthy individuals were recruited across two studies, the Alzheimer’s Disease Biomarkers (ADB) and Healthy Aging (HA) studies. Signed informed consent from all participants was obtained and the research protocol was approved by the Research Ethics Board of the Douglas Mental Health University Institute, Montreal, Canada. Exclusion criteria for both cohorts included history of neurological and psychiatric illness, physical injuries such as head trauma and concussion, alcohol/substance abuse or dependence, and current drug use. Data from these two cohorts were published in previous papers from our group (Bussy et al., 2020; Bussy et al., 2021; Tullo et al., 2019). The original data can be obtained through collaborative agreement and reasonable request but is not publicly available due to the lack of informed consent by these human participants. Complete demographic information of both samples is detailed in Table 1, and associated histograms for each variable are available in supplementary figure 1.

**Figure 1.**
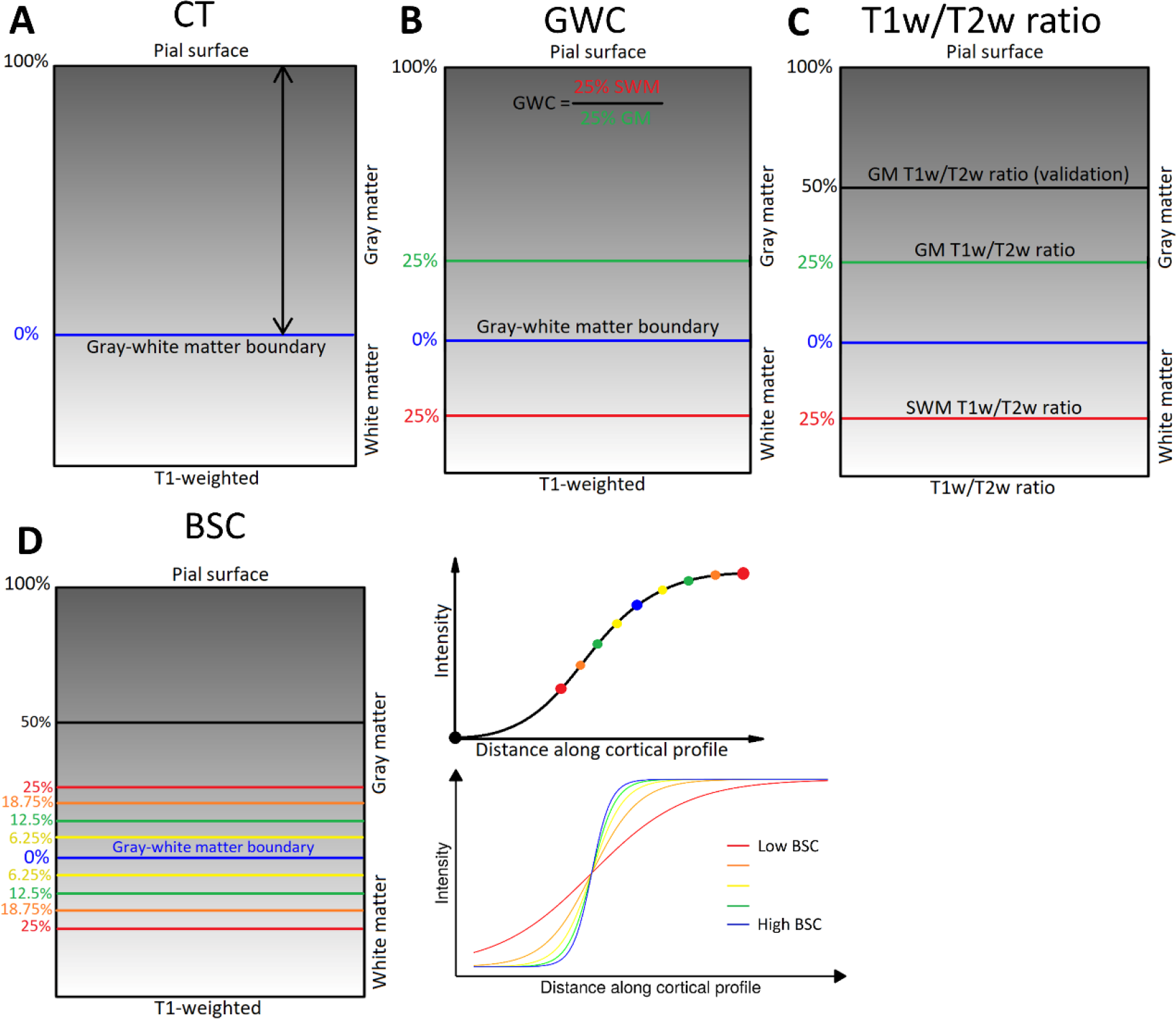
Methods for generating markers. **A.** Cortical Thickness (CT) estimates were calculated as the Laplacian distance between the pial surface and the gray-white matter boundary surface at each vertex on the T1-weighted volume in native space. **B.** Gray-white matter contrast (GWC) was calculated by dividing the intensity at 25% of CT translated into superficial white matter (SWM) by the intensity at 25% of CT into gray matter (GM) at each vertex on the T1-weighted volume in MNI space. **C.** The T1w/T2w ratio measures were generated by sampling the T1w/T2w volume in native space at various distances. GM T1w/T2w ratio was sampled at 25% of CT. SWM T1w/T2w ratio was sampled at 25% of CT translated into SWM. Additionally, a second GM T1w/T2w ratio measure was sampled at 50% of CT in GM (referred to as GM T1w/T2w ratio at 50% of CT). **D.** The boundary sharpness coefficient (BSC) was calculated by first sampling 10 T1-weighted intensities in MNI space around the gray-white matter boundary (between 50% of CT in GM and 25% of CT in SWM), then fitting a sigmoid curve to the resulting intensity profile at each vertex. The BSC represents the growth parameter of the sigmoid curve, with a higher BSC indicating a sharper gray-white matter transition and a lower BSC representing a more gradual transition.

**Table 1.** Subject demographics. Demographic information by dataset before quality control (QC), after QC and in the R1 subsample. HA = Healthy Aging cohort, ADB = Alzheimer’s Disease Biomarkers cohort (controls only)

|  | Pre-QC |  |  | Post-QC |  |  | R1 subsample |  |  |
| --- | --- | --- | --- | --- | --- | --- | --- | --- | --- |
|  | Total | ADB | HA | Total | ADB | HA | Total | ADB | HA |
| N | 174 | 68 | 106 | 127 | 45 | 82 | 35 | 26 | 9 |
| Mean age<br>(years +/- SD) | 55.3<br>+/-<br>17.59 | 69.9<br>+/-<br>5.59 | 45.8<br>+/-<br>16.12 | 53.4 +/-<br>17.97 | 69.6<br>+/-<br>5.72 | 44.5 +/-<br>16.1 | 62.7 +/-<br>13.33 | 70 +/-<br>5.17 | 41.8 +/-<br>4.87 |
| Sex<br>(female:male) | 95 : 79 | 41 : 27 | 54 : 52 | 76 : 51 | 29 :<br>16 | 47 : 35 | 21 : 14 | 16 :<br>10 | 5 : 4 |
| WASI (+/- SD) | 104.3<br>+/-<br>18.8 | - | 104.3<br>+/-<br>18.8 | 105.5<br>+/- 18.8 | - | 105.5<br>+/- 18.8 | 102.1<br>+/- 21.1 | - | 102.1<br>+/- 21.1 |
| MMSE (+/- SD) | 28.6<br>+/- 1.5 | 28.3<br>+/- 1.7 | 28.8<br>+/- 1.3 | 28.7 +/-<br>1.5 | 28.2<br>+/- 1.8 | 29 +/-<br>1.3 | 28.5 +/-<br>1.6 | 28.3<br>+/- 1.8 | 29 +/-<br>0.8 |
| MOCA (+/- SD) | 25.5<br>+/- 2.7 | 25.5<br>+/- 2.7 | - | 25.2 +/-<br>2.8 | 25.2<br>+/- 2.8 | - | 25.2 +/-<br>3 | 25.2<br>+/- 3 | - |
| RBANS (+/- SD) | 99.9<br>+/-<br>13.8 | 97.1<br>+/-<br>12.9 | 101.6<br>+/- 14 | 101.3<br>+/- 13 | 98.2<br>+/-<br>13.6 | 102.9<br>+/- 12.4 | 98 +/-<br>13 | 94.5<br>+/-<br>12.8 | 106.6<br>+/- 9.1 |

|  | Pre-QC |  |  | Post-QC |  |  | R1 subsample |  |  |
| --- | --- | --- | --- | --- | --- | --- | --- | --- | --- |
|  | Total | ADB | HA | Total | ADB | HA | Total | ADB | HA |
| N | 174 | 68 | 106 | 127 | 45 | 82 | 35 | 26 | 9 |
| Mean age<br>(years +/- SD) | 55.3<br>+/-<br>17.59 | 69.9<br>+/-<br>5.59 | 45.8<br>+/-<br>16.12 | 53.4 +/-<br>17.97 | 69.6<br>+/-<br>5.72 | 44.5 +/-<br>16.1 | 62.7 +/-<br>13.33 | 70 +/-<br>5.17 | 41.8 +/-<br>4.87 |
| Sex<br>(female:male) | 95 : 79 | 41 : 27 | 54 : 52 | 76 : 51 | 29 :<br>16 | 47 : 35 | 21 : 14 | 16 :<br>10 | 5 : 4 |
| WASI (+/- SD) | 104.3<br>+/-<br>18.8 | - | 104.3<br>+/-<br>18.8 | 105.5<br>+/- 18.8 | - | 105.5<br>+/- 18.8 | 102.1<br>+/- 21.1 | - | 102.1<br>+/- 21.1 |
| MMSE (+/-<br>SD) | 28.6<br>+/- 1.5 | 28.3<br>+/- 1.7 | 28.8<br>+/- 1.3 | 28.7 +/-<br>1.5 | 28.2<br>+/- 1.8 | 29 +/-<br>1.3 | 28.5 +/-<br>1.6 | 28.3<br>+/- 1.8 | 29 +/-<br>0.8 |
| MOCA (+/-<br>SD) | 25.5<br>+/- 2.7 | 25.5<br>+/- 2.7 | - | 25.2 +/-<br>2.8 | 25.2<br>+/- 2.8 | - | 25.2 +/-<br>3 | 25.2<br>+/- 3 | - |
| RBANS (+/-<br>SD) | 99.9<br>+/-<br>13.8 | 97.1<br>+/-<br>12.9 | 101.6<br>+/- 14 | 101.3<br>+/- 13 | 98.2<br>+/-<br>13.6 | 102.9<br>+/- 12.4 | 98 +/-<br>13 | 94.5<br>+/-<br>12.8 | 106.6<br>+/- 9.1 |

#### Alzheimer’s Disease Biomarkers (ADB)

In this cohort, subjects were recruited across the Alzheimer’s Disease spectrum, but only the healthy controls were included in this study (N = 68, 27 males and 41 females, mean age = 69.93 +/- 5.63, age range 56-81). The cognitive status of the participants was evaluated using two validated cognitive screening tests, namely the Mini-Mental State Exam (MMSE; Arevalo-Rodriguez et al., 2015) and Montreal Cognitive Assessment (MoCA; Nasreddine et al., 2005), and their cognition was further evaluated with the Repeatable Battery for the Assessment of Neuropsychological Status (RBANS; Randolph et al., 1998). Subjects with a MMSE ≥24/30 and a MoCA ≥26/30 were categorized in the control group.

#### Healthy Aging (HA)

In this cohort, participants throughout the healthy adult lifespan were recruited (N = 106, 54 females and 52 males, mean age = 45.37 +/- 16.20, age range 18-80). The cognitive abilities of the subjects were also evaluated with the MMSE and RBANS, and their IQ was assessed with the Wechsler Abbreviated Scale of Intelligence (WASI; Wechsler, 2007).

### 2.2 MRI acquisition

For all participants, T1w and T2w sequences were acquired at the Cerebral Imaging Center, associated with the Douglas Research Center in Montréal, Canada. Quantitative R1 (the inverse of the raw T1 signal, i.e. 1/T1) maps were acquired using the magnetization-prepared two rapid acquisition gradient echo (MP2RAGE) sequence on a subset of participants. All scans were conducted on the same Siemens Trio 3T MRI scanner using a 32-channel head coil.

#### T1-weighted

For both cohorts, the T1w magnetization-prepared rapid acquisition gradient echo (MPRAGE) sequence was acquired using parameters established by the Alzheimer’s Disease Neuroimaging Initiative (Jack et al., 2008); repetition time [TR] = 2300 ms; echo time [TE] = 2.98 ms; inversion time [TI] = 900 ms; flip angle [α] = 9°; GRAPPA = 2; slice thickness = 1 mm for 1 mm isotropic voxels and a total scan time of 5:12.

#### T2-weighted

T2w images were acquired using a SPACE sequence with the following parameters (TR = 2500 ms; TE = 198 ms; FOV = 206 mm; slice thickness = 0.64 mm for 0.64 mm isotropic voxel dimensions). The slice partial Fourier was set to 6/8 for the T2w scan of the ADB cohort, as a means to shorten the scan time and reduce the likelihood of motion artifacts. While this technique slightly decreases the signal-to-noise ratio, the image contrast should not be affected (Feinberg et al., 1986), and the two T2w sequences should be directly comparable. The total scan times for T2w images were 10:02 minutes for the Alzheimer’s Disease Biomarkers cohort and 13:16 minutes for the Healthy Aging cohort.

#### MP2RAGE

The MP2RAGE sequence (Marques et al., 2010) was acquired for a subset of the participants (N = 52) with the following parameters (TI1 = 700ms; TI2 = 2000ms, TE = 2.01ms; TR = 5000ms; α1 = 4°; α1 = 5°; FOV = 256 x 240 mm^2^; slice thickness = 0.8mm for 0.8mm isotropic voxels and a total scan time of 10:42).

### 2.3 Image processing

#### 2.3.1 Motion quality control

Exclusions were made based on a rigorous motion quality control (QC) procedure that was performed on all raw scans, following guidelines established by our laboratory (https://github.com/CoBrALab/documentation/wiki/Motion-Quality-Control-Manual) (Bedford et al., 2019). More specifically, T1w, T2w, and MP2RAGE scans were rated on a 4-point scale based on visible artifacts that are attributed to motion, such as ringing and blurring of the images. A higher score was indicative of poorer scan quality. Images that were ascribed a score above 2 were excluded from analyses, which is considered a strict criterion and has shown to provide robust estimates after downstream image processing (Bedford et al., 2019). As a result, 32 subjects that failed motion QC for either T1w or T2w images were excluded from the overall cohort, and 5 subjects that failed motion QC of the R1 maps were excluded from the R1 subsample.

#### 2.3.2 Preprocessing

T1w images were preprocessed using the minc-bpipe-library (https://github.com/CobraLab/minc-bpipe-library). The procedure consists of a N4 bias field correction (Tustison et al., 2010), cropping of the neck region, and brain extraction using the BEaST algorithm (Eskildsen et al., 2012). An example of the preprocessed T1w volume for one subject is available in supplementary figure 12. Since the processing of the T1w/T2w ratio requires native images (Glasser & Van Essen, 2011), no preprocessing steps were performed for the T2w images.

#### 2.3.3 Generation of cortical surfaces

From these outputs, brain volumes were transformed into standard MNI space using BestLinReg (Collins et al., 1994; Dadar et al., 2018), and the gray-white matter boundary and pial surfaces were generated with the CIVET 2.1.0 processing pipeline (https://www.bic.mni.mcgill.ca/ServicesSoftware/CIVET-2-1-0-References) (Kim et al., 2005).

The preprocessed T1w volumes were used for the extraction of CT, GWC and BSC measures.

#### 2.3.4 Surface quality control

The accuracy of the white matter and gray matter segmentations, as reflected in the surfaces generated by CIVET 2.1.0, was examined. Surfaces were quality controlled with a standardized procedure described in (Bedford et al., 2019) and https://github.com/CoBrALab/documentation/wiki/CIVET-Quality-Control-Guidelines. Scans were scored on a three-point scale based on the number and significance of segmentation errors, where a higher score was indicative of fewer segmentation errors. Examples of the most prominent artifacts are under-estimation of white matter in sensorimotor areas, under-segmentation of the temporal pole, and misclassification of ventricles as white or gray matter. Scans with a score below 1 were excluded from analyses (N=15).

After rigorous quality control procedures for both the raw scans and the white and gray matter segmentations, a total of 127 participants (51 males and 76 females, mean age = 53.35 +/- 18.04, age range 18 to 81) were included in the main analyses. The R1 subsample consisted of 35 participants (9 from the Healthy Aging cohort and 24 from the Alzheimer’s Disease Biomarkers cohort, 14 males and 21 females, mean age = 62.71 +/- 13.52, age range 36 to 79). Other demographic variables are available in Table 1 and histograms are available in supplementary figure 1.

### 2.4 Cortical marker generation

In order to extract the cortical markers, surfaces generated by CIVET were used.

Figure 1 illustrates how each marker is calculated. All markers were surface smoothed with a 20mm full-width half-maximum (FWHM) heat kernel and were projected onto a common cortical surface mesh (the ICBM 152 2009b sym model) to enable cross-subject comparisons. Additionally, main analyses are rerun on markers smoothed with a 5mm

FWHM heat kernel applied after regressing out curvature in order to assess if our results are robust to those parameter variations (see supplementary figures 8-11).

#### 2.4.1. Mean curvature

Since cortical markers previously described have been shown to correlate with the curvature of the cortical surfaces (Olafson et al., 2021; Shafee et al., 2015), we acquired curvature estimates of the gray-white matter surface with the CIVET 2.1.0 pipeline in order to residualize the markers against mean curvature. This process is done after the smoothing procedure for analyses in the main text, and before the smoothing procedure in supplementary analyses (see supplementary figures 8-11).

#### 2.4.2 Cortical thickness

CT estimates were generated with the CIVET 2.1.0 processing pipeline. More specifically, CT was defined as the Laplace distance between the gray-white matter boundary surface and the pial surface at each vertex (Figure 1A). These surfaces were subsequently used for the processing of the other markers, by providing a base from which other surfaces were generated in order to sample the intensities at various fractions of CT.

#### 2.4.3 Gray-white matter contrast

GWC measures are calculated on the T1w volume linearly transformed in MNI space by dividing the white matter intensity sampled at a distance equivalent to 25% of the cortical thickness in the direction of white matter by the gray matter intensity sampled at 25% of the cortical thickness along the normal of the surface at each vertex (Figure 1B). The GM sampling distance was chosen because previous studies show higher rates of myelination changes in childhood at around ¼ of the cortical depth (Whitaker et al., 2016), potentially indicating a higher sensitivity to aging, while the WM sampling distance was chosen in order to minimize partial volume effects.

#### 2.4.4 Boundary sharpness coefficient

The BSC is defined as the growth parameter of a sigmoid function fit to a depth profile of 10 intensity values along a path perpendicular to the gray-white matter boundary surface. It was developed to address certain limitations of the GWC, namely its reliance on the exact gray-white matter boundary placement that is sometimes unreliable and its widespread correlation with the curvature of the cortex (Olafson et al., 2021). Indeed, the BSC is theoretically less affected by the boundary placement as it quantifies the transition between gray and white matter continuously, and has been shown to correlate only in limited regions to cortical curvature (Olafson et al., 2021).

The computation of the BSC is explained in detail in Olafson and colleagues (2021) but is briefly covered here (Figure 1D). First, gray matter surfaces linearly transformed in MNI space were generated at increasing percentile fractions of CT from the white matter surface towards the pial surface (0%, 6.25%, 12.5%, 18.75%, 25%, 50%). Second, white matter surfaces were generated at the same percentile fractions as the gray matter surfaces, but in the direction of the white matter. However, the 50% white matter surface was omitted, since some vertices were located in the gray matter (crossing over into the opposing gyral bank), particularly in thin gyral crowns. Third, the intensity of the T1w image linearly transformed in MNI space was sampled at each vertex of the gray and white matter surfaces. Fourth, a sigmoid curve was fitted to the 10 sampled intensities at each vertex using a non-linear least squares estimator. From those curves, the BSC, which is the growth parameter of the sigmoid function, was extracted at each vertex. Hence, high BSC values represent a sharper transition between gray and white matter, while low BSC values represent a more gradual transition. The values were then log-transformed in order to ensure that the assumption of a normal distribution of the general linear model was respected.

#### 2.4.5 T1w/T2w ratio

The T1w/T2w method was developed by Glasser & Van Essen (2011) following the rationale that the T1w and T2w sequences are both sensitive to myelin, but in opposite directions (T1w signal being proportional to the quantity of myelin, while T2w signal being inversely proportional to the quantity of myelin). Hence, the ratio of those two images enhances the contrast-to-noise ratio of myelin (Glasser & Van Essen, 2011). Using this technique, the authors reported general agreement of myelin-based cortical parcellations between those identified in the T1w/T2w ratio images and previous histological findings (Glasser & Van Essen, 2011). However, while this technique has some histological support and reflects the quantity of myelin to some extent, it is more accurately a qualitative measure of myelin as the resulting signal is also influenced by molecule size, oligodendrocyte markers, mitochondria, and pH (Ritchie et al., 2018).

To generate the T1w/T2w ratio measures, it was first necessary to upsample the T1w images to 0.64 mm isotropic voxel dimensions (i.e. the same resolution as the T2w images) using a windowed sinc interpolation, as in Tullo and colleagues (2019). Then, to enable voxel-by-voxel correspondence between the T1w and T2w images in native space, T2w images were rigidly registered to T1w images using BestLinReg (Collins et al., 1994; Dadar et al., 2018). The two volumes were then mathematically divided to obtain the T1w/T2w ratio images. An example of the resulting volume for one subject is available in supplementary figure 12. The CIVET surfaces of each subject generated on T1w images were registered and transformed to the subject-specific T1w/T2w ratio volume.

Subsequently, the T1w/T2w ratio values were sampled at both 25% and 50% of CT in GM at each vertex (Figure 1C). This was in order to make sure that our results were not dependent on the specific cortical depth at which the T1w/T2w ratio was sampled. However, we chose the T1w/T2w ratio at 25% of CT to be the primary GM T1w/T2w ratio analysis, as this sampling distance is the same as the GM sampling distance of the GWC, and given the higher concentration of myelin at more superficial layers previously described (Whitaker et al., 2016). Furthermore, to obtain SWM T1w/T2w values, the T1w/T2w ratio was sampled at 25% of CT translated in the direction of white matter.

#### 2.4.6 R1

Quantitative MRI sequences differ from conventional (i.e., weighted) MRI sequences in that they directly measure the absolute relaxation times of the observable protons. As a result, they are a more interpretable measure of the physical properties of the tissue, and to some degree of the biology (Weiskopf et al., 2021). Since they are not influenced by extrinsic factors (e.g., acquisition parameters, specific hardware specifications, etc.), the images can theoretically be directly compared between scanners and studies (Deoni, 2010). The rate of longitudinal relaxation time R1 (1/T1) has been shown to be positively correlated with myelin content (Stüber et al., 2014). Since R1 is more specific to the underlying physical properties of the tissue than the T1w/T2w ratio, we assessed the extent of the spatial overlap between the two measures.

The R1 images (Marques et al., 2010) in native space were used to extract GM and SWM R1 values. The CIVET surfaces of each subject generated on T1w images were registered and transformed to the subject-specific R1 volume. The R1 values were sampled at both 25% and 50% of CT in GM at each vertex, and at 25% of CT in the direction of white matter. An example of the R1 volume for one subject is available in supplementary figure 12.

### 2.5 Examination of relationships with histological data

The “BigBrain” is an ultrahigh resolution (up to 20 um isotropic voxel dimension) digital reconstruction of a complete brain which was sliced and stained for cell bodies (Amunts et al., 2013) using the Merker staining method (Merker, 1983), where areas with high cellular density (e.g., gray matter) have a low intensity value and areas with low cellular density (e.g., white matter) have a high intensity value (i.e., intensity is inversely related to the density of cells), as can be seen in supplementary figure 12. The brain was donated by a 65 year old male. The digital reconstruction was then non-linearly registered to the standard MRI template ICBM152 (Fonov et al., 2009). High-resolution gray-white matter boundary and pial surfaces were then generated on the BigBrain volume downsampled to 400 um isotropic voxels in MNI space (Lewis et al., 2014). In this study, we used the 8-bit 400 um resolution BigBrain volume in MNI ICBM 152 space, which is the same space as our MRI results. It is important to note that the BigBrain surfaces were downsampled from 163842 vertices per hemisphere to 40962 vertices per hemisphere (i.e. the same number of vertices as MRI CIVET surfaces), thus allowing for direct comparison between BigBrain and MRI findings.

Markers on the BigBrain were generated in the same way as MRI markers, but on the BigBrain volume (Figure 1). CT estimates were calculated in native space. GWC and BSC measures were calculated as described above using the BigBrain volume and surfaces in MNI space. GM and SWM T1w/T2w ratio measures were compared with inverted BigBrain intensities sampled at the same distances (i.e. 25% of CT in GM, 50% of CT in GM, and 25% of CT in SWM). These values were inverted to simplify the interpretation of results since BigBrain intensities are inversely related to cell density. All sampled intensity values, used by all markers except for CT, were divided by 100 so that the value range was approximately in the same realm as MRI intensities (otherwise BSC and GWC values were very small and went beyond the computer’s numerical precision).

### 2.6 Statistical analyses

#### 2.6.1 Correlation with mean curvature

We observed widespread vertex-wise correlations between curvature and the BSC, GWC and CT metrics (see supplementary figure 2). As such, curvature was regressed out of all markers at the vertex-wise level to limit its influence on our downstream analyses for age-related trajectories.

**Figure 2.**
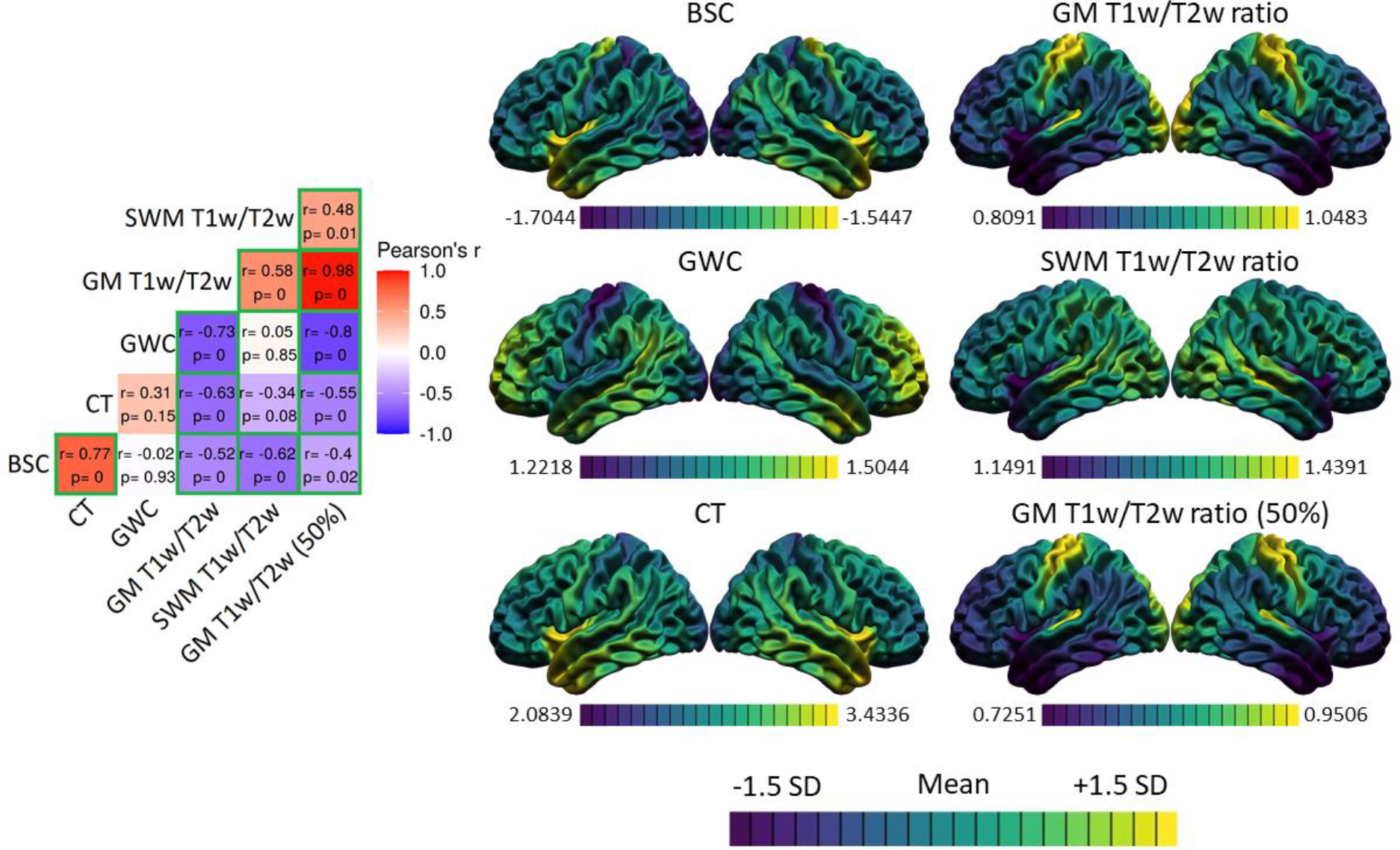
Spatial distributions of the markers and correlations. For each marker, the mean and standard deviation of the surface were calculated and used to threshold the colors. Purple areas indicate lower values relative to the mean of that marker, while yellow areas indicate higher values. The correlation matrix includes Pearson’s correlation coefficients (r) and FDR-corrected p-values. The color of each correlation block is linked to the correlation coefficient: positive coefficients are red and negative coefficients are blue, and high coefficients are more saturated and low coefficients tend towards white. Significant correlations at the FDR 0.05 level are highlighted with a green outline.

**Figure 3.**
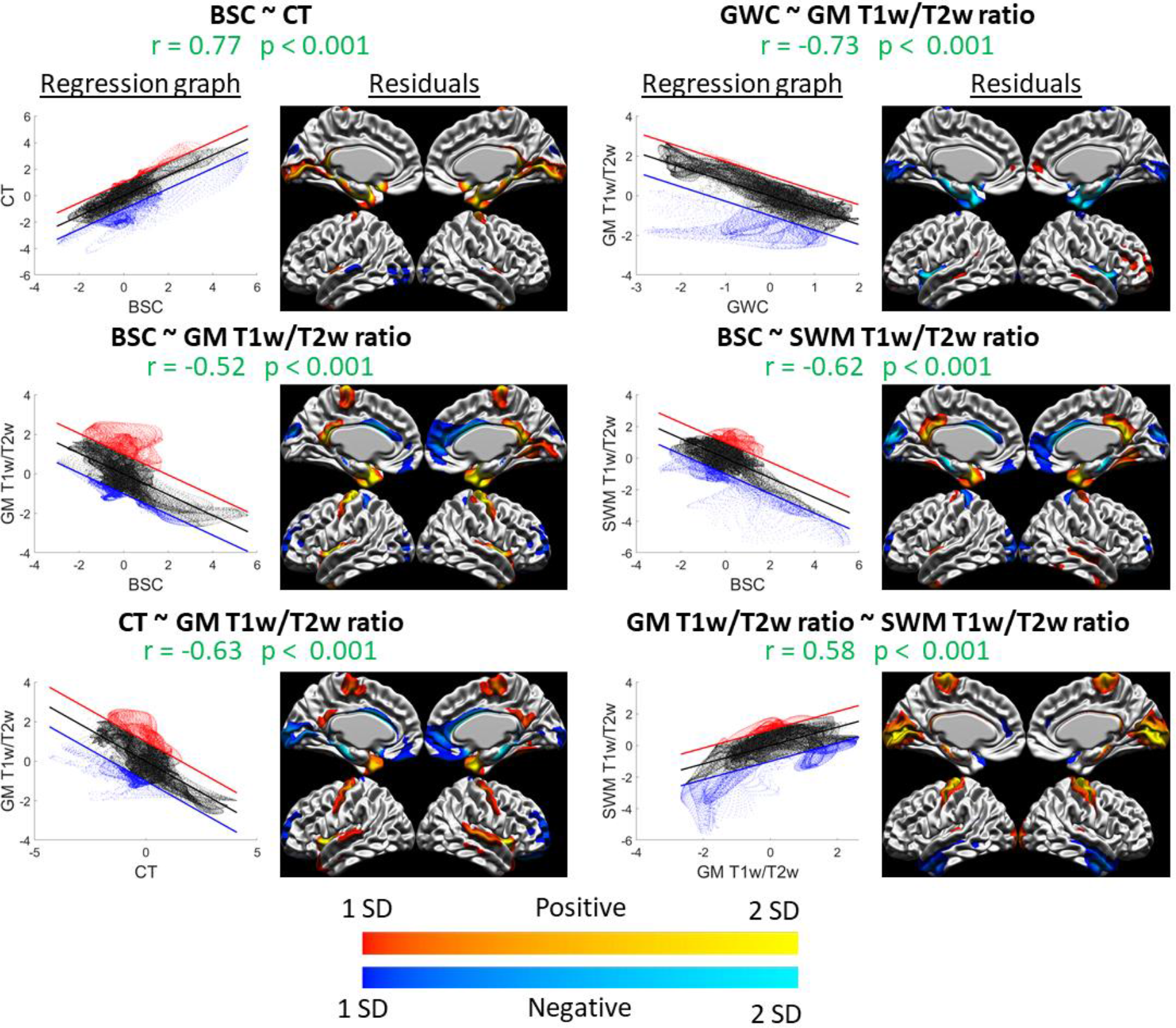
Spatial distribution relationships: graphs and residuals. For each significant correlation, the left figure is the spatial regression in graph form, where the x-axis are the values of the first marker, the y-axis are values of the second marker, the regression line is shown in black, and the +1 SD and -1 SD lines are shown in red and blue respectively. The right figure is the vertex-wise residuals from the regression thresholded at +/- 1 standard deviation (cold colors indicate vertices below the regression line in blue in the left graph and warm colors indicate vertices above the regression line in red in the left graph, and lighter colors indicate higher residual values and darker colors indicate lower residual values). For example, the relationship between GWC and GM T1w/T2w ratio (top right) is linear in most areas, as seen in the graph on the left, except for a group of vertices below the regression line which we can locate in the residual figure on the right (in this case, in the insula and medial temporal pole)

#### 2.6.2 Comparing the spatial distributions

The spatial distribution of each marker was generated by calculating the vertex-wise mean value across-subjects, which were then mapped to the cortical surface for comparison. The values used for this analysis were not residualized for curvature, since by definition, the sum of residuals following a least-squares fitting procedure always equals to 0, thus rendering the mean of residuals meaningless. The spatial correlations between the surface maps were hypothesis-tested following the “spin test” procedure detailed in see section 2.6.7.

#### 2.6.3 Comparing the shape of age trajectories

Age trajectories were modeled using linear models in R version 3.5.1 (https://www.r-project.org), more specifically with the vertexLm function of the RMINC package version 1.5.2.3 (Lerch et al., 2017). To evaluate the shape of the age trajectory of each marker, we compared linear, quadratic, and cubic models of age, with sex as a covariate, at each vertex using the Akaike information criterion (AIC) as in our previous work (Bedford et al., 2019; Bussy et al., 2021; Tullo et al., 2019):

1: Marker ∼ age + sex
2: Marker ∼ age + age^2^ + sex
3: Marker ∼ age + age^2^ + age^3^ + sex

The AIC respects the principle of parsimony by penalizing every additional predictor variable added to the statistical model (Mazerolle, 2006). Hence, the model with the lowest AIC at each vertex was considered the model which best fit the data. The results were mapped onto the common cortical surface for visualization and the number of vertices that were best fitted by each model was computed for each marker in order to compare the shape of the age trajectories between the markers (see Figure 4).

**Figure 4.**
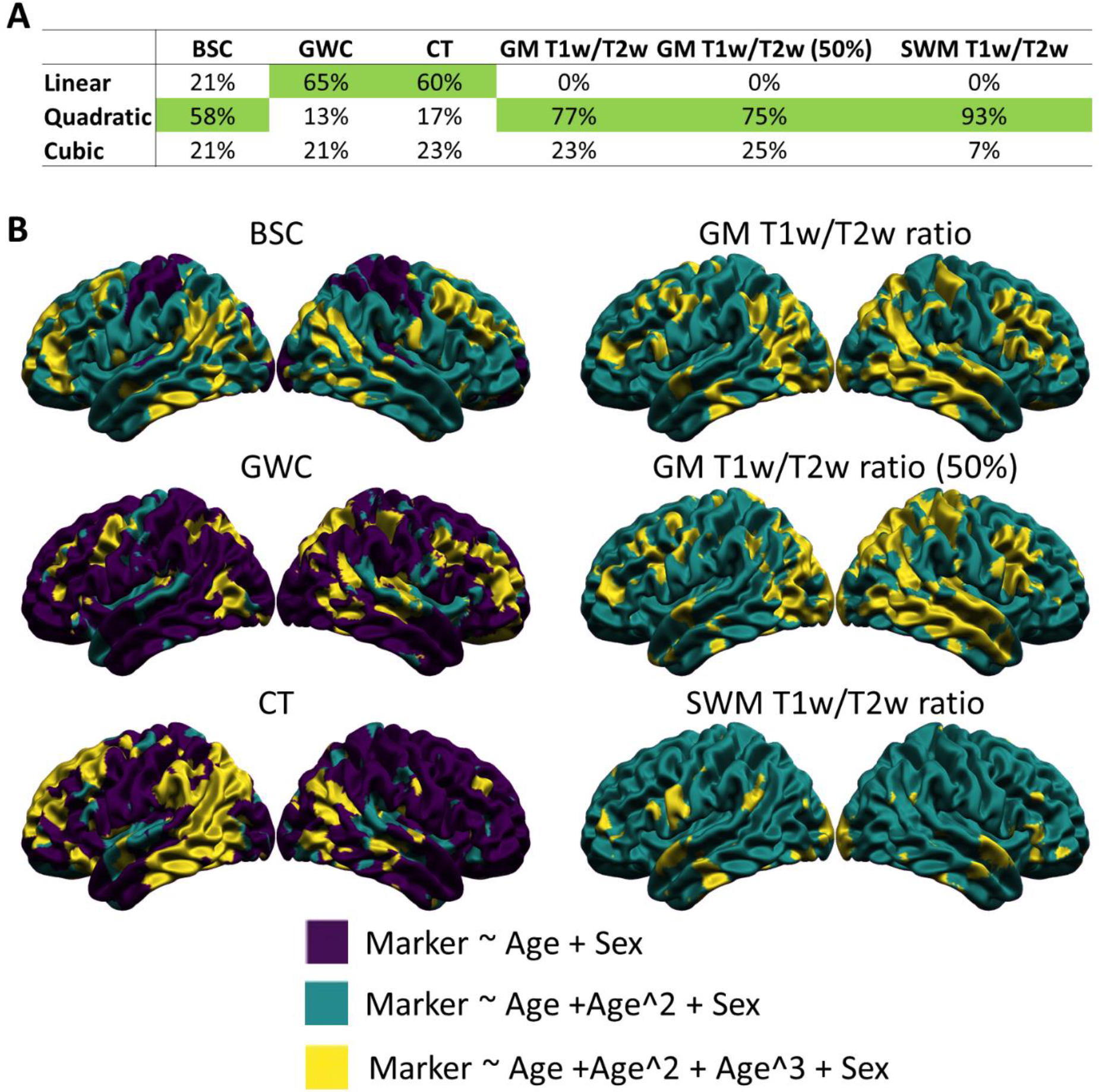
Vertex-wise best age trajectory shape between linear, quadratic and cubic for each marker. **A.** Table illustrating the proportion of vertices best fitted by each age model for each marker according to the Akaike Information Criterion (AIC), with the age model best fitting the highest proportion of vertices highlighted in green. **B.** Spatial distribution of the AIC results. Purple areas indicate a better fit of the linear age trajectory, green areas indicate a better fit of the quadratic age trajectory, and yellow areas indicate a better fit of the cubic age trajectory.

#### 2.6.4 Comparing the spatial distribution of the age trajectories

In order to further quantitatively compare the age trajectories between the markers, we compared the spatial distribution of the age effect. To do so, using a unified and simple age model for all markers was necessary. As a result, a linear age model with sex as a covariate (model 1 above) was fit at each vertex of each marker. The resulting p-values were corrected for multiple comparisons using the False Discovery Rate (FDR) correction, which controls the proportion of null hypotheses that are falsely rejected (Genovese et al., 2002).

The betas of the age component were then mapped onto the common cortical surface for comparison (see Figure 5A). The correlations between the surface maps were then assessed and hypothesis-tested following the “spin test” procedure detailed below in section 2.6.7. We visualized linear trajectories for each marker at a single vertex in the precentral gyrus, where the linear age effect of all markers was significant (see Figure 5B).

**Figure 5.**
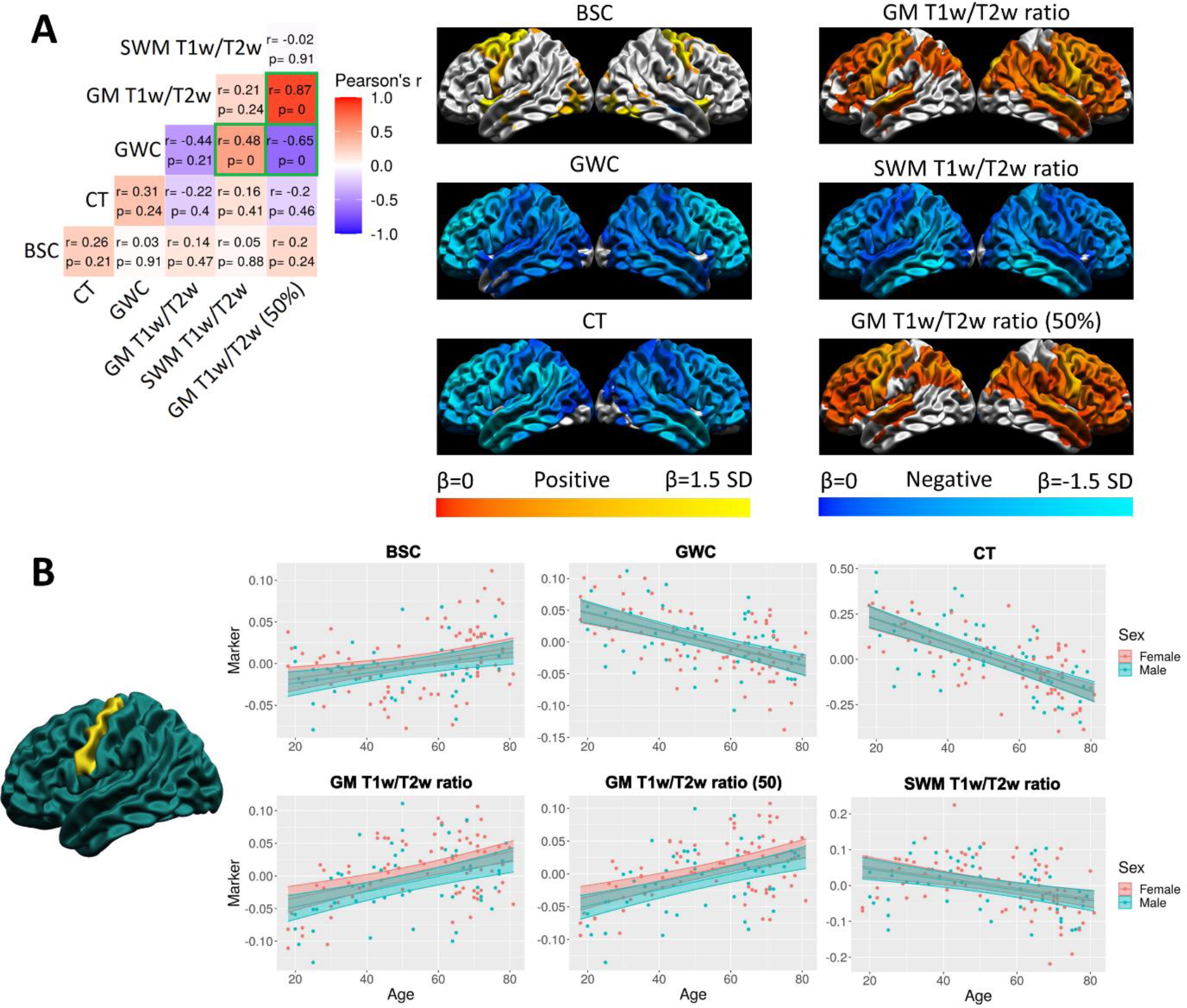
Spatial distribution of the linear age effect of the markers and correlations. **A.** For each marker, the mean and standard deviation of the age betas were calculated and used to threshold the colors. Cortical maps are thresholded for significance at the FDR 0.05 level. Cold colors indicate negative age betas and warm colors indicate positive age betas. Light colors indicate higher age betas and dark colors indicate lower age betas. The correlation matrix includes Pearson’s correlation coefficient (r) and FDR-corrected p-values. The color of each correlation block is linked to the correlation coefficient: positive coefficients are red and negative coefficients are blue, and high coefficients are more saturated and low coefficients tend towards white. Significant correlations at the FDR 0.05 level are highlighted with a green outline. **B.** Example of the age trajectory of each marker at one vertex in the precentral gyrus where the age beta of each marker was significant at the FDR 0.05 level. Blue observations represent male participants and red observations represent female participants. The x-axis is age and the y-axis is the marker value residualized for mean curvature.

#### 2.6.5 Comparing MRI and BigBrain markers

In order to assess the potential impact of cell density on the markers, the spatial correspondence between markers generated on MRI and on BigBrain was assessed (see Figure 6). More specifically, the same cortical maps of spatial distribution from MRI data, which were generated by calculating the vertex-wise across-subjects mean values of the markers, were used. While we cannot assume that the values are directly comparable between these two modalities because of the difference in intensity values, the spatial distribution of the values can be compared. Hence, the correlations between the surface maps were assessed and hypothesis-tested following the “spin test” procedure detailed below in section 2.6.7.

**Figure 6.**
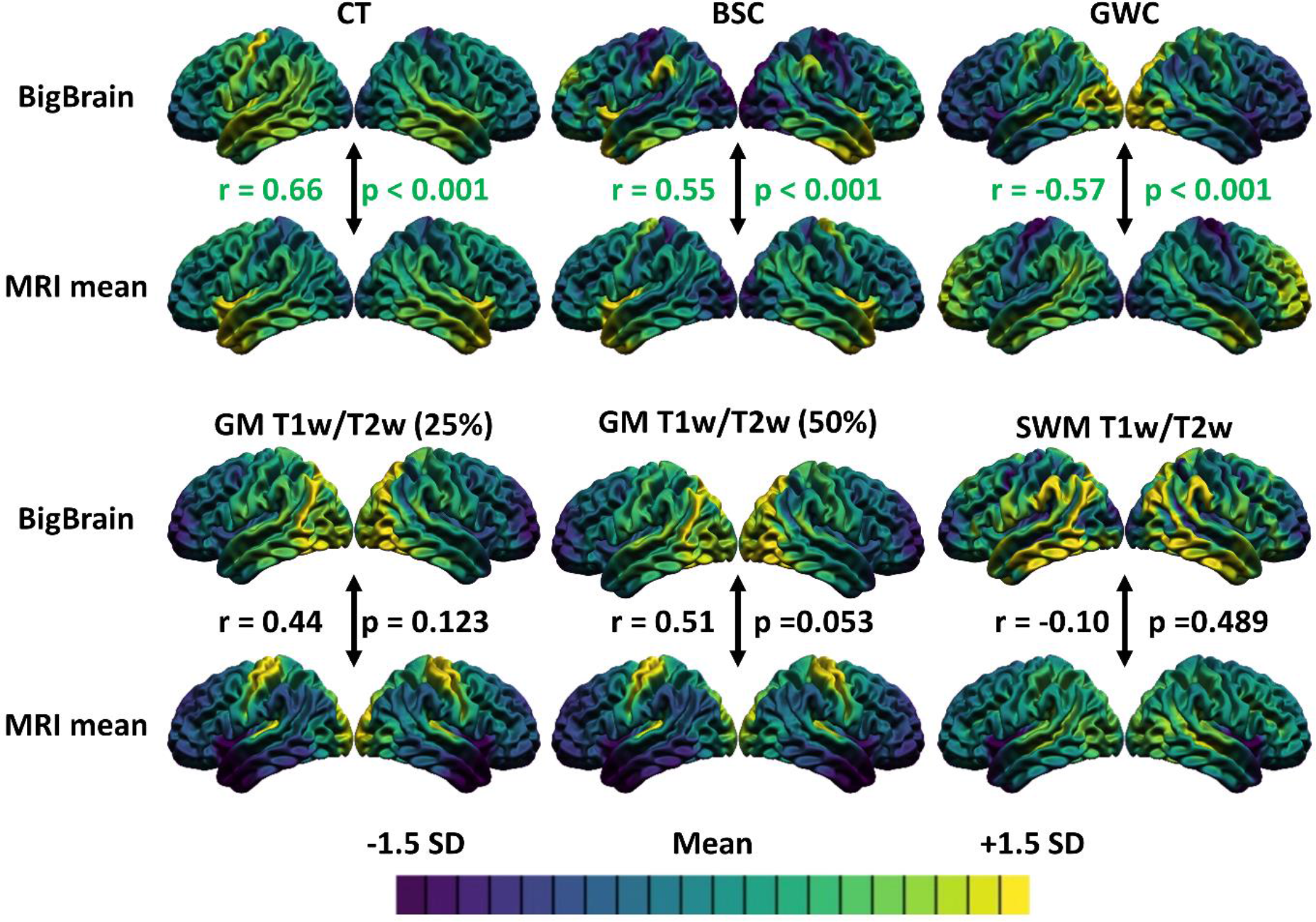
Correlations between the spatial distributions of the markers generated on MRI and markers generated on BigBrain. For each marker, the mean and standard deviation were calculated and used to threshold the colors. More specifically, purple areas indicate lower values relative to the mean of that marker, while yellow areas indicate higher values. The Pearson’s correlation coefficient and p-value are colored green if the relationship is significant at the 0.05 level.

#### 2.6.6 Comparing T1w/T2w ratio findings with R1

Using only the subjects in the subsample that passed QC of the R1 map, the spatial distribution correlations between BSC, GWC, CT, GM T1w/T2w ratio, SWM T1w/T2w ratio, GM R1 and SWM R1 were assessed (see supplementary figure 6). The correlations were then hypothesis-tested following the “spin test” procedure detailed below in section 2.6.7.

#### 2.6.7 Spatial correspondence between surface maps

The spatial correspondence between bilateral surface maps (i.e., left and right hemispheres bundled together) generated with the previous steps was assessed with the Pearson’s correlation coefficient. Each correlation was then hypothesis-tested using a bilateral ‘spin test’ (Alexander-Bloch et al., 2018). This novel statistical technique generates a null distribution of the spatial overlap by performing a large number of random rotations to spherical projections of the surfaces around each axis. Critically, this approach maintains the spatial relationship between vertices, in contrast with conventional parametric approaches that falsely assume independence of each vertex which lead to excessively high rates of false positives (Alexander-Bloch et al., 2018). In our analyses, we chose to do 1000 permutations, because results have been shown to converge between 500 and 1000 permutations (Markello & Misic, 2020). This technique outputs a p-value for each correlation. The p-values were then corrected for multiple comparisons using the FDR within each analysis (i.e. each correlation matrix) and considered significant below the 0.05 threshold. Of note, the medial wall was excluded from all analyses since the cortical markers are not valid in those regions. Furthermore, the correlation coefficient has to be interpreted in a spatial context. For example, if the correlation coefficient is positive, it means that where the values of the first surface are higher, the values on the second surface also tend to be higher, and vice versa.

In addition to the correlations, linear regressions on the Z-scored cortical maps were calculated. By definition, the beta of that regression, which is standardized, is exactly the same value as the Pearson’s correlation coefficient. This allows us to 1) graphically look at the relationship between cortical maps, and 2) map the residuals to the common cortical surface. It is then possible to assess which areas exhibit the relationship more (i.e. areas with lower residuals) and which areas exhibit the relationship less (i.e. areas with higher residuals).

## 3. Results

### 3.1 Correlation with mean curvature

The vertex-wise correlation of each marker with mean curvature was assessed (see supplementary figure 2). CT, GWC and BSC measures correlated significantly with mean curvature in regions across the cortex, while all T1w/T2w ratio measures only displayed a significant correlation in very small and isolated regions. However, significant correlations with mean curvature were less widespread for the BSC than for the GWC, supporting previous results (Olafson et al., 2021). In order for the markers to be directly comparable, all markers were regressed against mean curvature on a vertex-wise basis to limit the influence of curvature on the findings.

### 3.2 Comparing the spatial distributions

The vertex-wise mean values across-subjects were calculated for each marker (see Figure 2), resulting in metric-specific spatial distribution maps. The values used for this analysis were not residualized for curvature, since by definition, the sum of residuals following a least-squares fitting procedure always equals 0, thus rendering the mean of residuals meaningless. The values of the BSC were higher, indicating a sharper gray-white matter transition, in the temporal pole, the precentral gyrus, and the insula, while the values were lower in the occipital pole and postcentral regions, indicating a more gradual gray-white matter transition. For the GWC, the values were higher in lateral temporal regions, prefrontal lobe and temporo-parietal regions, while the values were lower in sensorimotor areas, occipital pole and insula. For CT, the values were higher in the temporal pole and insula, while the values were lower in the occipital lobe and postcentral gyrus. For the GM T1w/T2w ratio at both cortical depths (i.e. 25% of CT and 50% of CT), the values were higher in the occipital lobe and sensorimotor regions, while the values were lower in the frontal lobe, the lateral temporal and temporo-parietal regions. Lastly, for the SWM T1w/T2w ratio, the values were higher in the lateral occipital and superior temporal lobe while the values were lower in the insula and temporal pole.

The spatial correspondence between these surface maps was assessed with correlations, which were then hypothesis-tested via spin tests and the resulting p-values were corrected for multiple comparisons using the FDR (see Figure 2). There was a positive spatial correspondence between BSC and CT maps (r=0.77, p<0.001). Moreover, this relationship was exhibited across the vast majority of the cortex, as only a few regions in the medial temporal inferior cortex displayed high residuals (see Figure 3).

The spatial distribution of GM T1w/T2w ratio sampled at 25% of CT correlated negatively and significantly with all other markers, and the correlation was highest with the GWC (r=-0.73, p<0.001), followed by the correlation with CT (r=-0.63, p<0.001) and the correlation with the BSC (r=-0.52, p<0.001). In general, these relationships showed higher residuals in the precentral gyrus, the insula, and the lateral temporal pole, indicating a poorer fit of the correlation in those regions (see Figure 3). Also, a significant positive correlation was found between GM T1w/T2w ratio and SWM T1w/T2w ratio (r=0.58, p<0.001). This relationship showed higher residuals in sensorimotor regions, the medial occipital lobe, and the inferior lateral temporal lobe (see Figure 3). The correlations assessed with the GM T1w/T2w ratio at 50% of CT were not meaningfully different (see Figure 2).

Lastly, a significant and negative spatial correspondence was found between SWM T1w/T2w ratio and the BSC (r=-0.62, p<0.001), which showed higher residuals mostly in medial cortical regions (see Figure 3). All other correlations of the spatial distribution between the markers were not significant at the 0.05 level.

### 3.3 Comparing the shape of age trajectories

In order to determine which age trajectory shape was more appropriate for each marker, linear models with linear, quadratic, and cubic age variables at each vertex (with sex as a covariate) were compared using the AIC. For each marker, the number of vertices for which each age model was the best fit was counted (see Figure 4A). For CT and GWC measures, a linear model was the best fit for most vertices, followed by a cubic model and a quadratic model. For the BSC, a quadratic model was the best fit for most vertices, followed by a linear model and a cubic model. For all T1w/T2w ratio measures, a quadratic model was the best fit for the most vertices, followed by a cubic model and a linear model. The spatial distribution of age trajectory shapes is available in Figure 4B.

### 3.4 Comparing the spatial distribution of the age trajectories

To compare the spatial distribution of the age trajectories of the different markers, linear models with a linear age term and sex as a covariate were fit at each vertex. Since the goal of this study is not to best describe the age trajectories of the different markers, but to compare the age effect between markers, using a uniform and simpler linear age model across the markers is a preferred approach.

The betas of the age term were mapped onto the common cortical surface (see Figure 5A). A significant linear increase of the BSC with age was found primarily in anterior superior frontal regions, and in parts of the insula and lateral occipital lobe. For the GWC, a significant linear decrease with age was observed across most of the cortex, and this decrease was steeper in frontal regions. For CT, a similar widespread significant linear decrease with age was observed, which was steeper in frontal and temporal regions. For GM T1w/T2w ratio, a significant linear increase with age was found across most of the frontal lobe and in temporo-parietal regions, with the latter mostly in the right hemisphere, and no areas showed drastically steeper change with age. For SWM T1w/T2w ratio, a significant linear decrease was observed across most of the cortex, and was steeper in the inferior temporal lobe. Graphs of the linear age trajectories of each marker at one vertex in the precentral gyrus (where the linear age betas of all markers were significant) are available in Figure 5B.

The spatial correspondence between these surface maps was assessed with correlations, which were then hypothesis-tested via spin tests and the resulting p-values were corrected for multiple comparisons using the FDR (see Figure 5A). While the vertex-wise age betas shown in the figure are thresholded for significance at the FDR 0.05 level, the cortical maps that were correlated were not thresholded. There was a significant positive relationship between the spatial distribution of the age betas of the GWC and the SWM T1w/T2w ratio (r=0.48, p<0.001), meaning that where the GWC decreases more rapidly with age, the SWM T1w/T2w ratio also tends to decrease more rapidly. There was a significant negative relationship between the GWC and the GM T1w/T2w ratio only at mid cortical depth, meaning that where the GWC decreases more rapidly with age, the GM T1w/T2w ratio at mid cortical depth tends to increase more rapidly. Higher residuals of both relationships were mostly found in superior frontal and medial temporal regions (see supplementary figure 3).

Even if it is not the main goal of this article, the quadratic age trajectories of each marker were described, in order to compare our results with previous findings of quadratic age trajectories for some markers (Drakulich et al., 2021; Grydeland et al., 2013, 2019) and to accurately describe for the first time the quadratic age trajectory of the BSC. To do so, linear regressions with age linear, age quadratic and sex as predictors were fit at each vertex for each marker. The betas of the quadratic age term were then thresholded at the FDR 0.05 level and mapped to the common cortical surface (see supplementary figure 4A). For the BSC, the quadratic age term was significant and positive, indicating a u-shaped trajectory, across most of the cortex, except for sensorimotor regions and in the occipital pole. CT and GWC measures did not show significant quadratic age betas across the cortex. The T1w/T2w ratio, both in GM and in SWM, showed significant and negative quadratic age terms across the cortex, indicating inverted U-shaped age trajectories. Graphs of the quadratic age trajectories of each marker at one vertex in the precentral gyrus are available in supplementary figure 4B.

### 3.5 Comparing MRI and BigBrain markers

In order to probe the sensitivity of the markers to cellular organization, we compared the spatial distribution of the markers generated on BigBrain to the markers generated on MRI (see Figure 6). The spatial distribution of the MRI BSC was significantly positively correlated with the BigBrain BSC (r=0.55, p<0.001). Again, residuals were higher in the precentral gyrus and around the medial wall, although there were also high residuals sparsely distributed on the lateral cortical surface (see supplementary figure 5). For the GWC, there was a significant and negative correlation between the spatial distribution of the MRI GWC and BigBrain GWC (r=-0.57, p<0.001). Higher residuals of that relationship were observed in medial and lateral occipital lobe, insula, and medial temporal regions. For CT, there was a significant and positive correlation between the spatial distribution of MRI CT and BigBrain CT (r=0.66, p<0.001), while the residuals were higher around the medial wall, in the precentral gyrus, and in temporo-parietal regions. The correlation between the spatial distribution of GM BigBrain intensities and GM T1w/T2w ratio at 25% of CT was not significant (r=0.44, p=0.123), and neither was the correlation between SWM BigBrain intensities and SWM T1w/T2w ratio (r=-0.10, p=0.489). However, there was a positive spatial correlation approaching significance between GM BigBrain intensities at 50% of CT and GM T1w/T2w ratio at 50% of CT (r=0.51, p=0.053), with higher residuals in most of the lateral temporal lobe (except the temporal pole) and part of the prefrontal cortex.

### 3.6 Comparing the T1w/T2w ratio with R1 maps

The vertex-wise across-subjects mean values were calculated for each marker in the subsample of subjects that had a R1 map that passed quality control (see supplementary figure 6). Of note, the values used for this analysis were not residualized for curvature, as the sum of residuals from a linear regression following a least-squares fitting procedure is always equal to 0, thus rendering the mean meaningless. The spatial distribution of the BSC, GWC, CT, and T1w/T2w ratio measures were highly similar to the ones previously calculated in the whole sample. For the GM R1 at both cortical depths, the values were higher in the occipital lobe and sensorimotor regions, while the values were lower in the frontal lobe, the lateral temporal and temporo-parietal regions. For the SWM R1, the values were higher in the lateral occipital and superior temporal lobe while the values were lower in the insula and temporal pole.

The spatial correspondence between these surface maps was assessed with correlations, which were then hypothesis-tested via spin tests and the resulting p-values were corrected for multiple comparisons using the FDR (see supplementary figure 5). R1 and T1w/T2w ratio sampled at the same cortical depths showed very high positive spatial correspondence, which was slightly higher in GM (r=0.94, p<0.001) than in SWM (r=0.90, p<0.001). However, the small differences between R1 and T1w/T2w ratio resulted in bigger discrepancies in spatial correlations with other markers, at least in some instances. Indeed, the correlation between GM R1 and CT (r=-0.38, p=0.07) was lower than the correlation between GM T1w/T2w ratio and CT (r=-0.59, p<0.001). The correlation between SWM R1 and CT (r=-0.16, p=0.45) was lower than the correlation between SWM T1w/T2w ratio and CT (r=-0.35, p=0.06). The correlation between GM R1 and BSC (r=-0.36, p=0.05) was lower than the correlation between GM T1w/T2w ratio and BSC (r=-0.5, p<0.001). The correlation between SWM R1 and BSC (r=-0.44, p<0.001) was lower than the correlation between SWM T1w/T2w ratio and BSC (r=-0.63, p<0.001). However, other correlations with other markers were very similar between R1 and T1w/T2w ratio. Indeed, the correlation between GM R1 and GWC (r=-0.72, p<0.001) was the same as the correlation between GM T1w/T2w ratio and GWC (r=-0.72, p<0.001). The correlation between SWM R1 and GWC (r=-0.11, p=0.58) was highly similar to the correlation between SWM T1w/T2w ratio and GWC (r=0.00, p=0.99).

### 3.7 Influence of smoothing kernel and curvature regression order

The main analyses done on markers smoothed with a 20 mm FWHM heat kernel applied before regressing out curvature (Figures 2, 4-6) are compared with supplementary analyses done on markers smoothed with a 5 mm FWHM heat kernel applied after regressing out curvature (Supplementary figures 8-11). For the spatial distribution analysis, the supplementary analysis shows generally lower correlation coefficients, but a significant spatial correlation between SWM T1w/T2w ratio and CT not found in the main analysis. For the age trajectory shape analysis, the supplementary analysis shows a mainly linear age trajectory for the BSC, compared to a mainly quadratic age trajectory reported in the main analysis. Results also visually appear more noisy with the 5 mm smoothing kernel. For the spatial distribution of the linear age effect, significant linear changes with age are less spatially extensive and visually noisier in the supplementary analysis, but spatial correlations between markers are more significant (i.e., lower p-values), although correlation coefficients remain generally low and similar to ones in the main analysis. For the BigBrain-MRI comparisons, the supplementary analysis show generally lower spatial correlations between BigBrain-derived markers and MRI-derived markers, although the GM T1w/T2w ratio at 50% of CT and SWM T1w/T2w ratio correlations are significant as opposed to the main analysis. Again, in this analysis, cortical maps again appear visually more noisy in the supplementary analysis.

## 4. Discussion

In this study, we assessed the similarities and differences of commonly used MRI cortical markers that aim to quantify pericortical myelin and microstructure, namely CT, GWC, BSC, and T1w/T2w ratio. Although these measures are sometimes used interchangeably, there is significant relevance to compare these measures against one another and to assess their biological sensitivity to cytoarchitectural information derived from human histological data.

Our results show high correlations between the spatial distributions of these markers, indicating that the gross anatomical distribution of these markers could stem from the same microstructural property. However, the age trajectories of these markers diverge to a large extent, both in the shape, direction, and spatial distribution of the age effect, indicating that different microstructural properties are likely at the source of more subtle age-related changes.

### 4.1 Similarities and disparities with literature

At the level of individual metrics, our results are highly consistent with the literature. Indeed, we found spatial distributions in of CT (Fjell et al., 2009), GWC (Salat et al., 2009), BSC (Olafson et al., 2021), T1w/T2w ratio (Glasser & Van Essen, 2011), and R1 measures (Sereno et al., 2013) that correspond with previously reported spatial patterns. This provides confidence in our methodology and in the generalizability of our findings outside our sample.

Furthermore, we have replicated the age trajectories of the markers described in the literature. Indeed, a widespread linear decrease of CT in healthy aging was observed both in (Fjell et al., 2009) and in our sample, with a steeper age-related decline in frontal regions.

For the GWC, our observation of a linear decline with aging higher in the frontal lobe was consistent with previous findings (Vidal-Piñeiro et al., 2016). For the GM T1w/T2w ratio, we reproduce the well-known inverted-U-shaped age trajectory (Grydeland et al., 2013, 2019). To our knowledge, this is the first time the age trajectory of SWM T1w/T2w ratio was characterized, showing an inverted-U-shaped aging pattern similar to GM T1w/T2w ratio, but with a steeper decline in the elderly. This is somewhat similar to previously described age trajectories of fractional anisotropy (FA) in SWM (Nazeri et al., 2015), but we observed a more pronounced increase in SWM T1w/T2w ratio in early adulthood and a delayed decline compared to SWM FA. Compared to one study reporting a mixture of inverted-U shaped and linear age trajectories of the magnetization transfer ratio (MTR) in SWM, another measure sensitive to myelin, our SWM T1w/T2w ratio quadratic age trajectories are more widespread (Wu et al., 2016). Lastly, we have described the age effect of the BSC in the adult lifespan for the first time. Paired with the developmental trajectories described in (Olafson et al., 2021), we can describe for the first time the general age trajectory of the BSC across the whole lifespan: the boundary between GM and SWM becomes more gradual during childhood and adolescence, plateaus in adulthood, and becomes sharper in the elderly, showing a U-shaped age trend across the whole lifespan.

One discrepancy with the literature is the negligible correlation between T1w/T2w ratio and curvature observed in our sample compared to more extensive correlations reported in other studies (Shafee et al., 2015). Possible explanations for this discrepancy include a smaller sample size (1555 vs 127 participants in our study), different surface extraction method (FreeSurfer vs CIVET in our study), and smaller smoothing kernels (5mm vs 20mm FWHM kernels in our study). However, we replicate more widespread correlations with curvature for the GWC (Olafson et al., 2021) and CT (Sereno et al., 2013), and spatially restricted correlations for the BSC (Olafson et al., 2021).

### 4.2 Independence of GWC and BSC

An interesting finding was the observed independence of the BSC and the GWC, two measures that aim to represent similar cortical features, namely cortical blurring and contrast between gray and white matter. Indeed, the spatial distributions between BSC and GWC were completely uncorrelated (r=-0.02, p=0.93), the spatial distributions of the linear age effects were completely uncorrelated (r=0.03, p=0.91), and the age trajectory of the BSC showed a U-shaped age trajectory with increases in the elderly across the cortex while the age trajectory of the GWC showed a linear decline with age. This discrepancy between BSC and GWC has been observed in one study before (Olafson et al., 2021), and the authors have hypothesized the causes of this discrepancy to be differences in preprocessing, analyses methods, sample size and characteristics, quality control procedures, and the uncertainty of the boundary placement which would theoretically affect the GWC to a greater extent than the BSC. However, in the present study, we can exclude almost all of these possible biases, except for the boundary placement explanation, due to a rigorous matching between the two markers for processing, analyses, and sample.

One other explanation could be that the BSC values are driven by CT, possibly because the intensities of the cortical profile are sampled as fractions of CT. Indeed, there was a very high positive correlation between the spatial distributions of BSC and CT (r=0.77, p<0.001), which means that where BSC is higher, CT also tends to be higher. Hence, areas of high CT would induce deeper SWM and GM sampling distances, making the differences between SWM and GM intensities and possibly the sharpness of the GM/SWM boundary higher. However, it is also possible that CT is higher in areas of greater boundary sharpness, as the nature of correlations prevents us from determining the direction of the relationship.

On the other hand, while the BSC and CT show synchronous changes between childhood and adulthood (BSC and CT decrease), their trajectories diverge in the elderly, as the BSC gets higher while CT continues its linear decrease. Hence, it is possible that the gross anatomical distribution of the BSC is influenced by CT, but that smaller age-related changes are more independent. Determining the direction of the relationship between the BSC and CT would be highly relevant, as one marker could indicate a source of bias on the other.

In sum, it is possible that both BSC and GWC are valid measures that don’t capture the same cortical features. Indeed, it is possible that the contrast between gray and white matter becomes lower in aging, while the transition between the two entities becomes sharper.

### 4.3 Differences between spatial distributions and age trajectories

An important contribution of this paper is the observed discrepancy between spatial distribution relationships and aging relationships. Most of the correlations between the spatial distributions of the markers are significant at <0.05 FDR, and correlations range from r=-0.73 to r=0.77. As an example, GM T1w/T2w ratio, which we interpret as the most sensitive measure of intracortical myelin amongst all markers, correlated significantly and negatively with the BSC, GWC and CT, which could be interpreted as a myelin-specific dependency for these markers. On the other hand, the spatial correlations of the linear age effect between the markers are mostly non-significant at <0.05 FDR and range from r=-0.43 to r=0.48. Furthermore, the shape of the age trajectories differs, with the BSC showing quadratic U-shaped trajectories, T1w/T2w ratio measures showing quadratic inverted U-shaped trajectories, and CT and GWC showing linear decline trajectories. This discrepancy highlights an important point: microstructural properties driving spatial distributions of MRI markers can be different from microstructural properties driving, in our case, age-related effects. This rationale can also potentially extend to other pathology-related effects.

As an example, the spatial distribution of the BSC correlated negatively with the GM T1w/T2w ratio, meaning that where the BSC tends to be higher, the GM T1w/T2w ratio tends to be lower. Meanwhile, the spatial distribution of the linear age effect of the BSC does not correlate significantly with the GM T1w/T2w ratio. Hence, the microstructural properties impacting the spatial distribution of the BSC are different from the microstructural properties at the source of the age effect.

This finding warrants caution in the interpretation of age- and pathology-related MRI effects as being driven by the same microstructural properties as the spatial distribution. In other words, our finding of spatial correlations being higher for mean values than for linear age effect indicates that while some measures tend to covary at the cortex-wide level, age trajectories are likely influenced by different interactions of microstructural changes.

### 4.4 Similarity between T1w/T2w ratio and R1

We assessed the similarity between T1w/T2w ratio and R1 measures in the subsample of participants that had a clean R1 map (N = 35). The spatial distribution correlations between T1w/T2w ratio and R1 measures sampled at the same depths show very high similarities (between r=0.90 and r=0.94; see supplementary figure 6), which is in accordance with previous reports (Shams et al., 2019). However, relationships between these measures and BSC, GWC, and CT differed, with the spatial correlations being higher for T1w/T2w ratio than R1. This can possibly be explained by the fact that the BSC, GWC, and CT measures are generated on T1w images, which are also included in the calculation of the T1w/T2w ratio. This finding highlights that subtle differences between T1w/T2w ratio and R1 can lead to larger differences when correlating these measures to other measures, both in the results and interpretations (i.e. some relationships were significant at the <0.05 FDR only with T1w/T2w ratio and not R1). Also, considering that the R1 maps are more specific to myelin, our findings suggest that the most myelin-specific cortical marker derived from weighted MRI images is the T1w/T2w ratio given its higher correspondence with R1.

This is further supported by the qualitatively similar inverted-U shaped age trajectories between T1w/T2w ratio observed in this study and R1 reported in other studies (Erramuzpe et al. 2021). A more quantitative assessment of this relationship was not possible in this study due to the limited sample size of the R1 subsample, but would be highly pertinent.

### 4.5 Relationships with cytoarchitecture

We observed relatively few meaningful correlations between the MRI- and BigBrain-derived markers. First, while the spatial distribution of the MRI BSC did correlate significantly with the BigBrain BSC, it is highly likely this relationship is mediated by CT, given the very high positive correlation between the spatial distribution of the BSC and CT, and the high positive correlation between MRI CT and BigBrain CT. Hence, this finding is not informative of the possible influence of cytoarchitecture on the BSC.

Secondly, we found a significant negative correlation between the spatial distribution of the MRI GWC and BigBrain GWC. This is somewhat visually evident when looking at the sagittal medial view of the BigBrain and the T1w images (see supplementary figure 12): the GM/SWM contrast in cell density is higher in posterior and lower in anterior regions, which is contrary to the contrast T1w intensities. This relationship is likely due to the nature of the BigBrain dataset, where the intensities are inversely related to the cell density, thus rendering comparisons difficult. The relationship between cytoarchitecture and MRI signal is better expressed with the correlations between BigBrain intensities and T1w/T2w ratio measures.

Indeed, we found a positive correlation approaching significance between the spatial distribution of the GM T1w/T2w ratio and the inverted BigBrain intensities only at 50% of cortical depth and not at 25%. While not quite reaching significance, this increased dependence of the T1w signal on the density of cells in cortical areas of scarce myelination is in agreement with (Eickhoff et al., 2005). It is thus possible that the T1w/T2w ratio depends more on cytoarchitecture in more superficial layers of GM, but this causal relationship would need to be empirically tested. Interestingly, this relationship was less expressed in temporo-parietal regions and in parts of the prefrontal cortex, showing higher residuals from the correlation. Higher residuals in those areas are either due to the BigBrain donor’s brain being different than the group average in these regions, or due to a different relationship between MRI signal and cytoarchitecture in these regions than the rest of the cortex. In partial support of the first hypothesis, the residuals from the BigBrain-MRI relationship of CT, a metric less influenced by modality differences, showed partial overlap with the regions mentioned above, mostly in temporo-parietal regions.

In sum, further investigations are needed to assess the impact of cytoarchitecture on the MRI cortical markers. Despite significant correlations between the BigBrain- and MRI-derived BSC and GWC, and correlations approaching significance for the T1w/T2w ratio only at half cortical depth, we cannot conclude with confidence that these markers are impacted to a large extent by cytoarchitecture. On the other hand, the influence of cytoarchitecture on weighted MRI images is not to be discarded, especially in more superficial cortical layers.

### 4.6 High residuals in medial cortex

One incidental finding in our analyses was the high residuals from most spatial correlations in medial regions, most prominently in the medial temporal lobe. While it is possible that this area has a different relationship between biological microstructure and MRI signal than the rest of the cortex, a more probable explanation is the unreliability of the measures caused by very low cortical thickness (<2mm), which could induce partial volume effects. Indeed, the GWC, BSC and T1w/T2w ratio measures are calculated by sampling the signal at different fractions of CT. Hence, very low CT could lead to the multiple sampling points being very close to each other, thus leading to unreliable measures. In order to test this hypothesis, we spatially correlated CT with the residuals from the 6 significant spatial distribution correlations (see supplementary figure 7B). Supporting this hypothesis, 4 out of 6 of those correlations were significant. Furthermore, we calculated the sum of squared residuals from the sigmoid curve used to generate the BSC at each vertex, then averaged the values across-subjects, and found high residuals in the same areas, meaning a worse fit of the sigmoid curve to the cortical profile in those areas (supplementary figure 6A).

Following those observations, we advise caution in interpreting results of these cortical markers in medial areas of very low CT.

### 4.7. Influence of smoothing kernel and curvature regression order

The main analyses done on markers smoothed with a 20 mm FWHM heat kernel applied before regressing out curvature (Figures 2, 4-6) are compared with supplementary analyses done on markers smoothed with a 5 mm FWHM heat kernel applied after regressing out curvature (Supplementary figures 8-11). In general, results from the supplementary analyses are similar and lead to similar conclusions compared to results from the main analyses, with the possible exception of a mostly linear age trajectory of the BSC in the supplementary analysis as opposed to a mostly quadratic age trajectory in the main analysis. Since cortical maps in supplementary analyses appear visually noisier, probably due to the lower smoothing kernel, we conclude that the higher smoothing kernel used in the main analyses was the most appropriate for our data. Interestingly, correlation coefficients are in general lower in the supplementary analyses, but p-values derived from spin tests (Alexander-Bloch et al., 2018) are generally lower (more significant). One possible explanation is that correlation coefficients from spinned surfaces (i.e. correlations forming the null distribution for the spin test) are disproportionately lower than the correlation of the non-spinned surfaces at lower smoothing kernel values, thus leading to a bigger difference between the original correlation and the null distribution resulting in a lower p-value. This interaction between p-values derived from spin-tests and smoothing kernels should be further investigated in the future.

### 4.8 Limitations

As a first limitation, it is important to note that our analyses are at the cortex-wide level. Hence, some interactions in local areas between markers and microstructure could still be present. For instance, (Natu et al., 2019) reported that the increased myelination of the cortex during development directly leads to reductions in CT specifically in the ventral temporal cortex, and this finding was validated histologically.

Secondly, our analyses cannot exclude that different characteristics of the myeloarchitecture could cause the age-related changes of the markers. In other words, the overall density of myelin in GM can be uncorrelated with some markers, but specific changes in laminar patterns of myelin could still differentially affect each of the markers, leading to dissimilar age trajectories that are caused by changes in myelination. For example, the GWC could represent the myelination similarity between GM and SWM, the BSC could represent the sharpness of myelin change between GM and SWM, and the T1w/T2w ratio could represent the density of myelin at the sampled cortical depth. However, we argue that such interpretation would need to be precisely characterized and empirically justified. It is also possible that the gross anatomical distribution of the markers, and of the T1w signal, represents myelin to a large extent, but that more subtle age- or disease-related changes could stem from changes in other microstructural properties also contributing to the signal, such as iron (Callaghan et al., 2014).

Thirdly, the linear age effects we used to compare the spatial distribution of the age trajectories between the markers are not optimal models in all cases, since the BSC and T1w/T2w ratio measures display mostly quadratic trajectories. However, those markers still display a significant linear component of those age trajectories, with the possible exception of the BSC showing somewhat spatially constrained significant linear age effects in anterior frontal areas. Furthermore, we argue that the different age trajectory shapes between the markers, rendering the quantitative comparison of the spatial distribution of the age effect more difficult, supports our interpretation that the age trajectories are different and driven by divergent microstructural changes.

Lastly, the observed correlations between the spatial distributions of MRI- and BigBrain-derived markers could be due to the idiosyncrasies of the BigBrain, which stems from a single brain of a 65-year-old male. However, it is unlikely that important differences between spatial distributions would arise from using different brains, hence our conclusions are unlikely to change.

### 4.9 Future work

Our results advise against attributing a specific microstructural property at the source of age- or pathology-related changes of MRI cortical biomarkers. While our findings highlight a discrepancy is the microstructural interpretation of the cortical markers, our conclusions do not aim to discourage the use of these MRI-derived markers, as they have been reported to be sensitive to various pathologies (Olafson et al., 2021; Salat et al., 2011), and could be useful for such purposes. In that regard, our finding of relative independence of the markers in aging indicates that they could be used complementarily as they could be sensitive to different cortical pathologies, thus highlighting the richness of information available in standard T1w and T2w images. Future work aiming to assess specific cortical microstructural properties should consider the use of multimodal quantitative MRI. Indeed, the advent of quantitative MRI allows for the unprecedented assessment of brain microstructural properties in-vivo, sometimes referred to as in-vivo histology (Weiskopf et al., 2021). Those techniques show increased biological specificity and are less sensitive to scanner- and sequence-specific differences, rendering them theoretically directly comparable between sites and scanners. However, the same rationale displayed here could also apply to quantitative MRI, meaning that the microstructural properties contributing the most to the contrast could be different from microstructural properties at the source of statistical effects. Hence, we advise for the use of multimodal quantitative MRI in order to increase the confidence of biological interpretations, as we have demonstrated in recent work from our group (Patel et al., 2019; Robert et al., 2021). However, as the adoption of quantitative MRI is lagging, our findings illustrate that many largely independent markers can be derived from the growing number of publicly available standard T1w and T2w scans, although specific biological interpretations of these markers would need to be further investigated.

### 4.10 Conclusion

In this study, we examined and compared the spatial distributions and age trajectories of the BSC, GWC, CT, and T1w/T2w ratio. These markers are all thought to be influenced by GM myelin (i.e. intracortical myelin), but evidence supporting these interpretations is lacking. Our results show similar spatial distributions between the markers, but few relationships in aging. Hence, we conclude that the microstructural properties at the source of spatial distributions of MRI cortical markers (e.g. GM myelin) can be different from microstructural changes that affect these markers in aging. This warrants care in interpreting the age- or disease-related effects of these MRI markers when aiming to show changes in a specific property of the cortical microstructure.

## Statements

### Author contributions

**Olivier Parent**: Conceptualization - Formal Analysis - Funding Acquisition - Investigation - Methodology - Software - Visualization - Writing Original Draft - Writing Review & Editing. **Emily Olafson**: Methodology - Visualization - Writing Review & Editing. **Aurélie Bussy**: Formal Analysis - Methodology - Writing Review & Editing. **Stephanie Tullo**: Resources - Writing Review & Editing. **Nadia Blostein**: Methodology - Software - Writing Review & Editing. **Alyssa Salaciak**: Resources - Writing Review & Editing. **Saashi A. Bedford**: Resources - Writing Review & Editing. **Sarah Farzin**: Resources - Writing Review & Editing. **Marie-Lise Béland**: Resources - Writing Review & Editing. **Vanessa Valiquette**: Data Curation - Resources - Writing Review & Editing. **Christine L. Tardif**: Conceptualization - Supervision - Writing Review & Editing. **Gabriel A. Devenyi**: Formal Analysis - Methodology - Software - Visualization - Writing Review & Editing. **M. Mallar Chakravarty**: Conceptualization - Formal Analysis - Funding Acquisition - Investigation - Methodology - Project Administration - Supervision - Writing Original Draft - Writing Review & Editing.

### Data and code availability statement

Original scans from participants can be obtained through collaborative agreement and reasonable request but are not publicly available due to the lack of informed consent by these human participants. However, we are committed to provide all anonymized single-subject brain maps used as inputs for analyses on the Canadian Open Neuroscience Platform. We will also provide the code used for these analyses and our obtained results in raw form on a public GitHub repository. All pipelines used for processing the MRI data are also publicly available tools, referenced throughout the manuscript.

### Ethics statement

Signed informed consent from all participants was obtained and the research protocol was approved by the Research Ethics Board of the Douglas Mental Health University Institute, Montreal, Canada.

## Acknowledgments

Parent is funded by the Fonds de Recherche du Québec Santé. M. Chakravarty is funded by the Weston Brain Institute, the Canadian Institutes of Health Research, the Natural Sciences and Engineering Research Council of Canada, the Fondation de Recherches Santé Québec and Healthy Brains for Healthy Lives.

## References

Alexander-Bloch, A. F., Shou, H., Liu, S., Satterthwaite, T. D., Glahn, D. C., Shinohara, R. T., Vandekar, S. N., & Raznahan, A. (2018). On testing for spatial correspondence between maps of human brain structure and function. NeuroImage, 178, 540–551. https://doi.org/10.1016/j.neuroimage.2018.05.070

Amunts, K., Lepage, C., Borgeat, L., Mohlberg, H., Dickscheid, T., Rousseau, M.-É., Bludau, S., Bazin, P.-L., Lewis, L. B., Oros-Peusquens, A.-M., Shah, N. J., Lippert, T., Zilles, K., & Evans, A. C. (2013). BigBrain: an ultrahigh-resolution 3D human brain model. Science, 340(6139), 1472–1475. https://doi.org/10.1126/science.1235381

Arevalo-Rodriguez, I., Smailagic, N., Roqué I Figuls, M., Ciapponi, A., Sanchez-Perez, E., Giannakou, A., Pedraza, O. L., Bonfill Cosp, X., & Cullum, S. (2015). Mini-Mental State Examination (MMSE) for the detection of Alzheimer’s disease and other dementias in people with mild cognitive impairment (MCI). Cochrane Database of Systematic Reviews. https://doi.org/10.1002/14651858.cd010783.pub2

Avino, T. A., & Hutsler, J. J. (2010). Abnormal cell patterning at the cortical gray–white matter boundary in autism spectrum disorders. Brain Research, 1360, 138–146. https://doi.org/10.1016/j.brainres.2010.08.091

Bedford, S. A., Park, M. T. M., Devenyi, G. A., Tullo, S., Germann, J., Patel, R., Anagnostou, E., Baron-Cohen, S., Bullmore, E. T., Chura, L. R., Craig, M. C., Ecker, C., Floris, D. L., Holt, R. J., Lenroot, R., Lerch, J. P., Lombardo, M. V., Murphy, D. G. M., Raznahan, A., … Chakravarty, M. M. (2019). Large-scale analyses of the relationship between sex, age and intelligence quotient heterogeneity and cortical morphometry in autism spectrum disorder. Molecular Psychiatry, 25(3), 614–628, https://doi.org/10.1038/s41380-019-0420-6

Bock, N. A., Kocharyan, A., Liu, J. V., & Silva, A. C. (2009). Visualizing the entire cortical myelination pattern in marmosets with magnetic resonance imaging. Journal of Neuroscience Methods, 185(1), 15–22. https://doi.org/10.1016/j.jneumeth.2009.08.022

Bussy, A., Patel, R., Plitman, E., Tullo, S., Salaciak, A., Bedford, S. A., Farzin, S., Béland, M.L., Valiquette, V., Kazazian, C., Tardif, C. L., Devenyi, G. A., & Chakravarty, M. M. (2021). Hippocampal shape across the healthy lifespan and its relationship with cognition. Neurobiology of Aging, 106, 153–168. https://doi.org/10.1016/j.neurobiolaging.2021.03.018

Bussy, A., Plitman, E., Patel, R., Salaciak, A., Farzin, S., Bedford, S., Béland, M., Tullo, S., Devenyi, G. A., & Chakravarty, M. M. (2020). Volumetric, shape and microstructural alterations of the hippocampal subfields in healthy aging. Alzheimer’s & Dementia, 16(4). https://doi.org/10.1002/alz.039589

Bussy, A., Plitman, E., Patel, R., Tullo, S., Salaciak, A., Bedford, S. A., Farzin, S., Béland, M.L., Valiquette, V., Kazazian, C., Tardif, C. L., Devenyi, G. A., & Chakravarty, M. M. (2021). Hippocampal subfield volumes across the healthy lifespan and the effects of MR sequence on estimates. NeuroImage, 233, 117931. https://doi.org/10.1016/j.neuroimage.2021.117931

Callaghan, M. F., Freund, P., Draganski, B., Anderson, E., Cappelletti, M., Chowdhury, R., Diedrichsen, J., FitzGerald, T. H. B., Smittenaar, P., Helms, G., Lutti, A., & Weiskopf, N. (2014). Widespread age-related differences in the human brain microstructure revealed by quantitative magnetic resonance imaging. Neurobiology of Aging, 35(8), 1862–1872. https://doi.org/10.1016/j.neurobiolaging.2014.02.008

Chwa, W. J., Tishler, T. A., Raymond, C., Tran, C., Anwar, F., Villablanca, J. P., Ventura, J., Subotnik, K. L., Nuechterlein, K. H., & Ellingson, B. M. (2020). Association between cortical volume and gray-white matter contrast with second generation antipsychotic medication exposure in first episode male schizophrenia patients. Schizophrenia Research, 222, 397–410. https://doi.org/10.1016/j.schres.2020.03.073

Collins, D. L., Neelin, P., Peters, T. M., & Evans, A. C. (1994). Automatic 3D intersubject registration of MR volumetric data in standardized Talairach space. Journal of Computer Assisted Tomography, 18(2), 192–205.

Dadar, M., Fonov, V. S., Collins, D. L., & Alzheimer’s Disease Neuroimaging Initiative. (2018). A comparison of publicly available linear MRI stereotaxic registration techniques. NeuroImage, 174, 191–200. https://doi.org/10.1016/j.neuroimage.2018.03.025

Deoni, S. C. L. (2010). Quantitative Relaxometry of the Brain. Topics in Magnetic Resonance Imaging: TMRI, 21(2), 101–113. https://doi.org/10.1097/rmr.0b013e31821e56d8

Drakulich, S., Thiffault, A. C., Olafson, E., Parent, O., Labbe, A., Albaugh, M. D., Khundrakpam, B., Ducharme, S., Evans, A., Chakravarty, M. M., & Karama, S. (2021). Maturational trajectories of pericortical contrast in typical brain development. NeuroImage, 235, 117974. https://doi.org/10.1016/j.neuroimage.2021.117974

Eickhoff, S., Walters, N. B., Schleicher, A., Kril, J., Egan, G. F., Zilles, K., Watson, J. D. G., & Amunts, K. (2005). High-resolution MRI reflects myeloarchitecture and cytoarchitecture of human cerebral cortex. Human Brain Mapping, 24(3), 206–215. https://doi.org/10.1002/hbm.20082

Erramuzpe, A., Schurr, R., Yeatman, J. D., Gotlib, I. H., Sacchet, M. D., Travis, K. E., Feldman, H. M., & Mezer, A. A. (2021). A Comparison of Quantitative R1 and Cortical Thickness in Identifying Age, Lifespan Dynamics, and Disease States of the Human Cortex. Cerebral Cortex, 31(2), 1211–1226. https://doi.org/10.1093/cercor/bhaa288

Eskildsen, S. F., Coupé, P., Fonov, V., Manjón, J. V., Leung, K. K., Guizard, N., Wassef, S. N., Østergaard, L. R., & Collins, D. L. (2012). BEaST: Brain extraction based on nonlocal segmentation technique. NeuroImage, 59(3), 2362–2373. https://doi.org/10.1016/j.neuroimage.2011.09.012

Feinberg, D. A., Hale, J. D., Watts, J. C., Kaufman, L., & Mark, A. (1986). Halving MR imaging time by conjugation: demonstration at 3.5 kG. Radiology, 161(2), 527–531. https://doi.org/10.1148/radiology.161.2.3763926

Fjell, A. M., Westlye, L. T., Amlien, I., Espeseth, T., Reinvang, I., Raz, N., Agartz, I., Salat, D. H., Greve, D. N., Fischl, B., Dale, A. M., & Walhovd, K. B. (2009). High Consistency of Regional Cortical Thinning in Aging across Multiple Samples. Cerebral Cortex, 19(9), 2001–2012. https://doi.org/10.1093/cercor/bhn232

Fonov, V. S., Evans, A. C., McKinstry, R. C., Almli, C. R., & Collins, D. L. (2009). Unbiased nonlinear average age-appropriate brain templates from birth to adulthood. NeuroImage, 47(S102). https://doi.org/10.1016/S1053-8119(09)70884-5

Fukunaga, M., Li, T.-Q., Van Gelderen, P., De Zwart, J. A., Shmueli, K., Yao, B., Lee, J., Maric, D., Aronova, M. A., Zhang, G., Leapman, R. D., Schenck, J. F., Merkle, H., & Duyn, J. H. (2010). Layer-specific variation of iron content in cerebral cortex as a source of MRI contrast. Proceedings of the National Academy of Sciences, 107(8), 3834–3839. https://doi.org/10.1073/pnas.0911177107

Genovese, C. R., Lazar, N. A., & Nichols, T. (2002). Thresholding of Statistical Maps in Functional Neuroimaging Using the False Discovery Rate. NeuroImage, 15(4), 870– 878. https://doi.org/10.1006/nimg.2001.1037

Glasser, M. F., & Van Essen, D. C. (2011). Mapping Human Cortical Areas In Vivo Based on Myelin Content as Revealed by T1- and T2-Weighted MRI. Journal of Neuroscience, 31(32), 11597–11616. https://doi.org/10.1523/jneurosci.2180-11.2011

Grydeland, H., Vértes, P. E., Váša, F., Romero-Garcia, R., Whitaker, K., Alexander-Bloch, A. F., Bjørnerud, A., Patel, A. X., Sederevičius, D., Tamnes, C. K., Westlye, L. T., White, S. R., Walhovd, K. B., Fjell, A. M., & Bullmore, E. T. (2019). Waves of Maturation and Senescence in Micro-structural MRI Markers of Human Cortical Myelination over the Lifespan. Cerebral Cortex, 29(3), 1369–1381. https://doi.org/10.1093/cercor/bhy330

Grydeland, H., Westlye, L. T., Walhovd, K. B., & Fjell, A. M. (2013). Improved prediction of Alzheimer’s disease with longitudinal white matter/gray matter contrast changes. Human Brain Mapping, 34(11), 2775–2785. https://doi.org/10.1002/hbm.22103

Jack, C. R., Jr, Bernstein, M. A., Fox, N. C., Thompson, P., Alexander, G., Harvey, D., Borowski, B., Britson, P. J., L Whitwell, J., Ward, C., Dale, A. M., Felmlee, J. P., Gunter, J. L., Hill, D. L. G., Killiany, R., Schuff, N., Fox-Bosetti, S., Lin, C., Studholme, C., … Weiner, M. W. (2008). The Alzheimer’s Disease Neuroimaging Initiative (ADNI): MRI methods. Journal of Magnetic Resonance Imaging: JMRI, 27(4), 685–691. https://doi.org/10.1002/jmri.21049

Jørgensen, K. N., Nerland, S., Norbom, L. B., Doan, N. T., Nesvåg, R., Mørch-Johnsen, L., Haukvik, U. K., Melle, I., Andreassen, O. A., Westlye, L. T., & Agartz, I. (2016). Increased MRI-based cortical grey/white-matter contrast in sensory and motor regions in schizophrenia and bipolar disorder. Psychological Medicine, 46(9), 1971–1985. https://doi.org/10.1017/s0033291716000593

Kim, J. S., Singh, V., Lee, J. K., Lerch, J., Ad-Dab’bagh, Y., MacDonald, D., Lee, J. M., Kim, S. I., & Evans, A. C. (2005). Automated 3-D extraction and evaluation of the inner and outer cortical surfaces using a Laplacian map and partial volume effect classification. NeuroImage, 27(1), 210–221. https://doi.org/10.1016/j.neuroimage.2005.03.036

Lerch, J., Hammill, C., van Eede, M., & Cassel, D. (2017). RMINC: Statistical Tools for Medical Imaging NetCDF (MINC) Files. R Package Version 1.5.2.1. http://mouse-imaging-centre.github.io/RMINC

Lewis, L.B., Lepage, C., Fournier, M., Zilles, K., Amunts, K., & Evans, A.C. (2014). BigBrain: Initial tissue classification and surface extraction. Annual Meeting of the Organization for Human Brain Mapping, Hamburg. https://bigbrain.loris.ca/papers/HBM2014poster.pdf

Markello, R. D., & Misic, B. (2021). Comparing spatially-constrained null models for parcellated brain maps. Neuroimage, 236, 118052. https://doi.org/10.1016/j.neuroimage.2021.118052

Marques, J. P., Kober, T., Krueger, G., van der Zwaag, W., Van de Moortele, P.F., & Gruetter, R. (2010). MP2RAGE, a self bias-field corrected sequence for improved segmentation and T1-mapping at high field. NeuroImage, 49(2), 1271–1281. https://doi.org/10.1016/j.neuroimage.2009.10.002

Mazerolle, M. (2006). Improving data analysis in herpetology: using Akaike’s Information Criterion (AIC) to assess the strength of biological hypotheses. Amphibia-Reptilia, 27(2), 169–180. https://doi.org/10.1163/156853806777239922

Merker, B. (1983). Silver staining of cell bodies by means of physical development. Journal of Neuroscience Methods, 9(3), 235–241. https://doi.org/10.1016/0165-0270(83)90086-9

Nasreddine, Z. S., Phillips, N. A., Bédiirian, V., Charbonneau, S., Whitehead, V., Collin, I., Cummings, J. L., & Chertkow, H. (2005). The Montreal Cognitive Assessment, MoCA: A Brief Screening Tool For Mild Cognitive Impairment. Journal of the American Geriatrics Society, 53(4), 695–699. https://doi.org/10.1111/j.1532-5415.2005.53221.x

Natu, V. S., Gomez, J., Barnett, M., Jeska, B., Kirilina, E., Jaeger, C., Zhen, Z., Cox, S., Weiner, K. S., Weiskopf, N., & Grill-Spector, K. (2019). Apparent thinning of human visual cortex during childhood is associated with myelination. Proceedings of the National Academy of Sciences, 116(41), 20750–20759. https://doi.org/10.1073/pnas.1904931116

Nazeri, A., Chakravarty, M. M., Rajji, T. K., Felsky, D., Rotenberg, D. J., Mason, M., Xu, L. N., Lobaugh, N. J., Mulsant, B. H., & Voineskos, A. N. (2015). Superficial white matter as a novel substrate of age-related cognitive decline. Neurobiology of Aging, 36(6), 2094–2106. https://doi.org/10.1016/j.neurobiolaging.2015.02.022

Olafson, E., Bedford, S. A., Devenyi, G. A., Patel, R., Tullo, S., Park, M. T. M., Parent, O., Anagnostou, E., Baron-Cohen, S., Bullmore, E. T., Chura, L. R., Craig, M. C., Ecker, C., Floris, D. L., Holt, R. J., Lenroot, R., Lerch, J. P., Lombardo, M. V., Murphy, D. G. M., … Chakravarty, M. M. (2021). Examining the Boundary Sharpness Coefficient as an Index of Cortical Microstructure in Autism Spectrum Disorder. Cerebral Cortex, 31(7), 3338–3352. https://doi.org/10.1093/cercor/bhab015

Patel, R., Steele, C. J., Chen, A. G. X., Patel, S., Devenyi, G. A., Germann, J., Tardif, C. L., & Chakravarty, M. M. (2019). Investigating microstructural variation in the human hippocampus using non-negative matrix factorization. NeuroImage, 207, 116348 . https://doi.org/10.1016/j.neuroimage.2019.116348

Patel, Y., Shin, J., Drakesmith, M., Evans, J., Pausova, Z., & Paus, T. (2020). Virtual histology of multi-modal magnetic resonance imaging of cerebral cortex in young men. NeuroImage, 218, 116968. https://doi.org/10.1016/j.neuroimage.2020.116968

Randolph, C., Tierney, M. C., Mohr, E., & Chase, T. N. (1998). The Repeatable Battery for the Assessment of Neuropsychological Status (RBANS): preliminary clinical validity. Journal of Clinical and Experimental Neuropsychology, 20(3), 310–319. https://doi.org/10.1076/jcen.20.3.310.823

Ritchie, J., Pantazatos, S. P., & French, L. (2018). Transcriptomic characterization of MRI contrast with focus on the T1-w/T2-w ratio in the cerebral cortex. NeuroImage, 174, 504–517. https://doi.org/10.1016/j.neuroimage.2018.03.027

Robert, C., Patel, R., Blostein, N., Steele, C., & Chakravarty, M. M. (2021). Microstructural variation in the human striatum using non-negative matrix factorization. *bioRxiv*. https://www.biorxiv.org/content/10.1101/2021.06.10.447764v1.abstract

Salat, D. H., Chen, J. J., Van Der Kouwe, A. J., Greve, D. N., Fischl, B., & Rosas, H. D. (2011). Hippocampal degeneration is associated with temporal and limbic gray matter/white matter tissue contrast in Alzheimer’s disease. NeuroImage, 54(3), 1795– 1802. https://doi.org/10.1016/j.neuroimage.2010.10.034

Salat, D. H., Lee, S. Y., Van Der Kouwe, A. J., Greve, D. N., Fischl, B., & Rosas, H. D. (2009). Age-associated alterations in cortical gray and white matter signal intensity and gray to white matter contrast. NeuroImage, 48(1), 21–28. https://doi.org/10.1016/j.neuroimage.2009.06.074

Sereno, M. I., Lutti, A., Weiskopf, N., & Dick, F. (2013). Mapping the Human Cortical Surface by Combining Quantitative T1 with Retinotopy†. Cerebral Cortex, 23(9), 2261–2268. https://doi.org/10.1093/cercor/bhs213

Shafee, R., Buckner, R. L., & Fischl, B. (2015). Gray matter myelination of 1555 human brains using partial volume corrected MRI images. NeuroImage, 105, 473–485. https://doi.org/10.1016/j.neuroimage.2014.10.054

Shams, Z., Norris, D. G., & Marques, J. P. (2019). A comparison of in vivo MRI based cortical myelin mapping using T1w/T2w and R1 mapping at 3T. PloS One, 14(7), e0218089. https://doi.org/10.1371/journal.pone.0218089

Stüber, C., Morawski, M., Schäfer, A., Labadie, C., Wähnert, M., Leuze, C., Streicher, M., Barapatre, N., Reimann, K., Geyer, S., Spemann, D., & Turner, R. (2014). Myelin and iron concentration in the human brain: A quantitative study of MRI contrast. NeuroImage, 93, 95–106. https://doi.org/10.1016/j.neuroimage.2014.02.026

Tardif, C. L., Gauthier, C. J., Steele, C. J., Bazin, P.-L., Schäfer, A., Schaefer, A., Turner, R., & Villringer, A. (2016). Advanced MRI techniques to improve our understanding of experience-induced neuroplasticity. NeuroImage, 131, 55–72. https://doi.org/10.1016/j.neuroimage.2015.08.047

Tullo, S., Patel, R., Devenyi, G. A., Salaciak, A., Bedford, S. A., Farzin, S., Wlodarski, N., Tardif, C. L., Breitner, J. C. S., Chakravarty, M. M., & the PREVENT-AD Research Group. (2019). MR-based age-related effects on the striatum, globus pallidus, and thalamus in healthy individuals across the adult lifespan. In Human Brain Mapping. https://doi.org/10.1002/hbm.24771

Tustison, N. J., Avants, B. B., Cook, P. A., Yuanjie, Z., Egan, A., Yushkevich, P. A., & Gee, J. C. (2010). N4ITK: Improved N3 Bias Correction. IEEE Transactions on Medical Imaging, 29(6), 1310–1320. https://doi.org/10.1109/tmi.2010.2046908

Vidal-Piñeiro, D., Walhovd, K. B., Storsve, A. B., Grydeland, H., Rohani, D. A., & Fjell, A. M. (2016). Accelerated longitudinal gray/white matter contrast decline in aging in lightly myelinated cortical regions. Human Brain Mapping, 37(10), 3669–3684. https://doi.org/10.1002/hbm.23267

Wechsler, D. (2007). Wechsler abbreviated scale of intelligence (WASI). Norwegian manual supplement. Stockholm, *Sweden*: *Pearson Assessment*.

Weiskopf, N., Edwards, L. J., Helms, G., Mohammadi, S., & Kirilina, E. (2021). Quantitative magnetic resonance imaging of brain anatomy and in vivo histology. Nature Reviews Physics. https://doi.org/10.1038/s42254-021-00326-1

Whitaker, K. J., Vértes, P. E., Romero-Garcia, R., Váša, F., Moutoussis, M., Prabhu, G., Weiskopf, N., Callaghan, M. F., Wagstyl, K., Rittman, T., Tait, R., Ooi, C., Suckling, J., Inkster, B., Fonagy, P., Dolan, R. J., Jones, P. B., Goodyer, I. M., & Bullmore, E. T. (2016). Adolescence is associated with genomically patterned consolidation of the hubs of the human brain connectome. Proceedings of the National Academy of Sciences, 113(32), 9105–9110. https://doi.org/10.1073/pnas.1601745113

Wu, M., Kumar, A., & Yang, S. (2016). Development and aging of superficial white matter myelin from young adulthood to old age: Mapping by vertex-based surface statistics (VBSS). Human Brain Mapping, 37(5), 1759–1769. https://doi.org/10.1002/hbm.23134

