## Supplementary figures for "High spatial overlap but diverging age-related trajectories of cortical MRI markers aiming to represent intracortical myelin and microstructure"

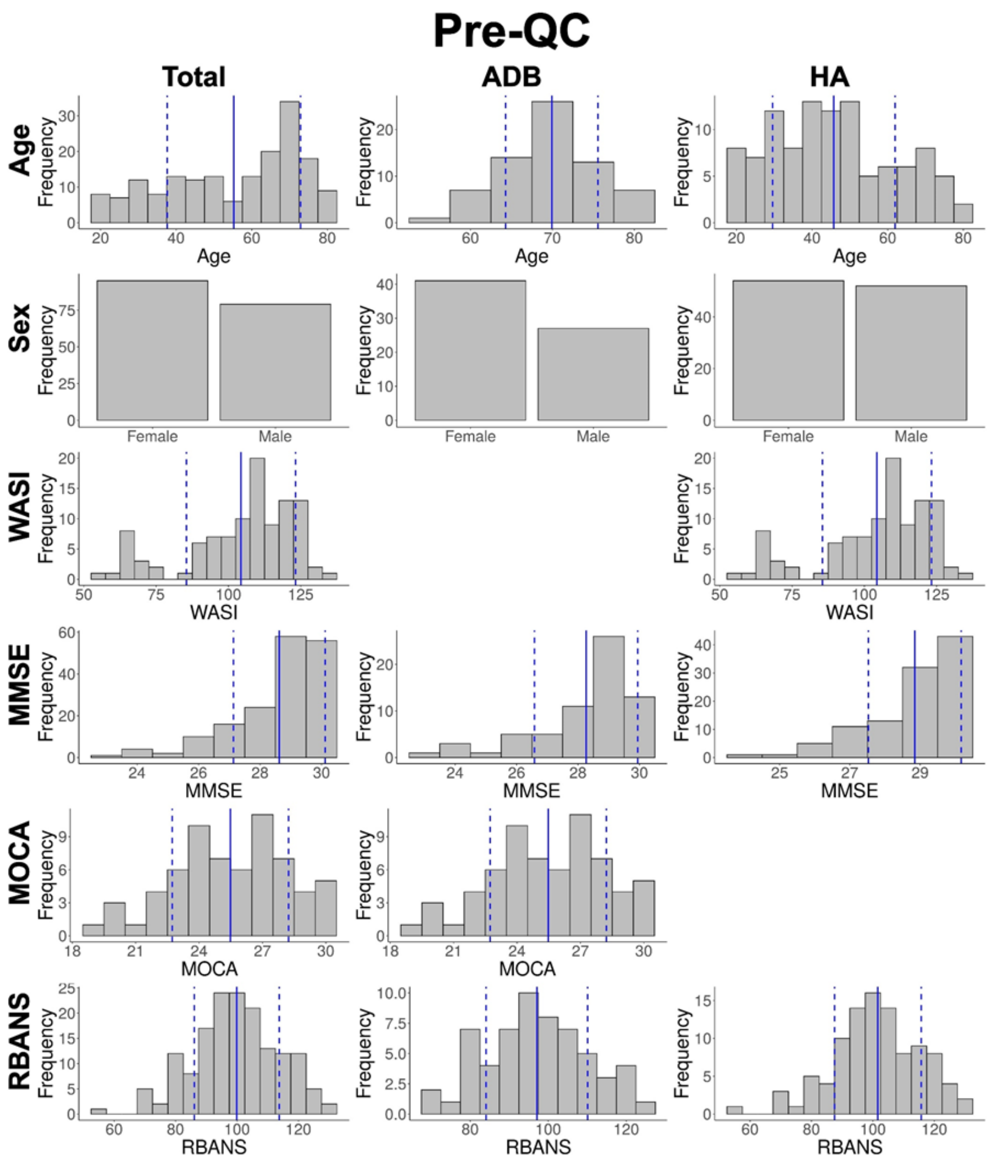

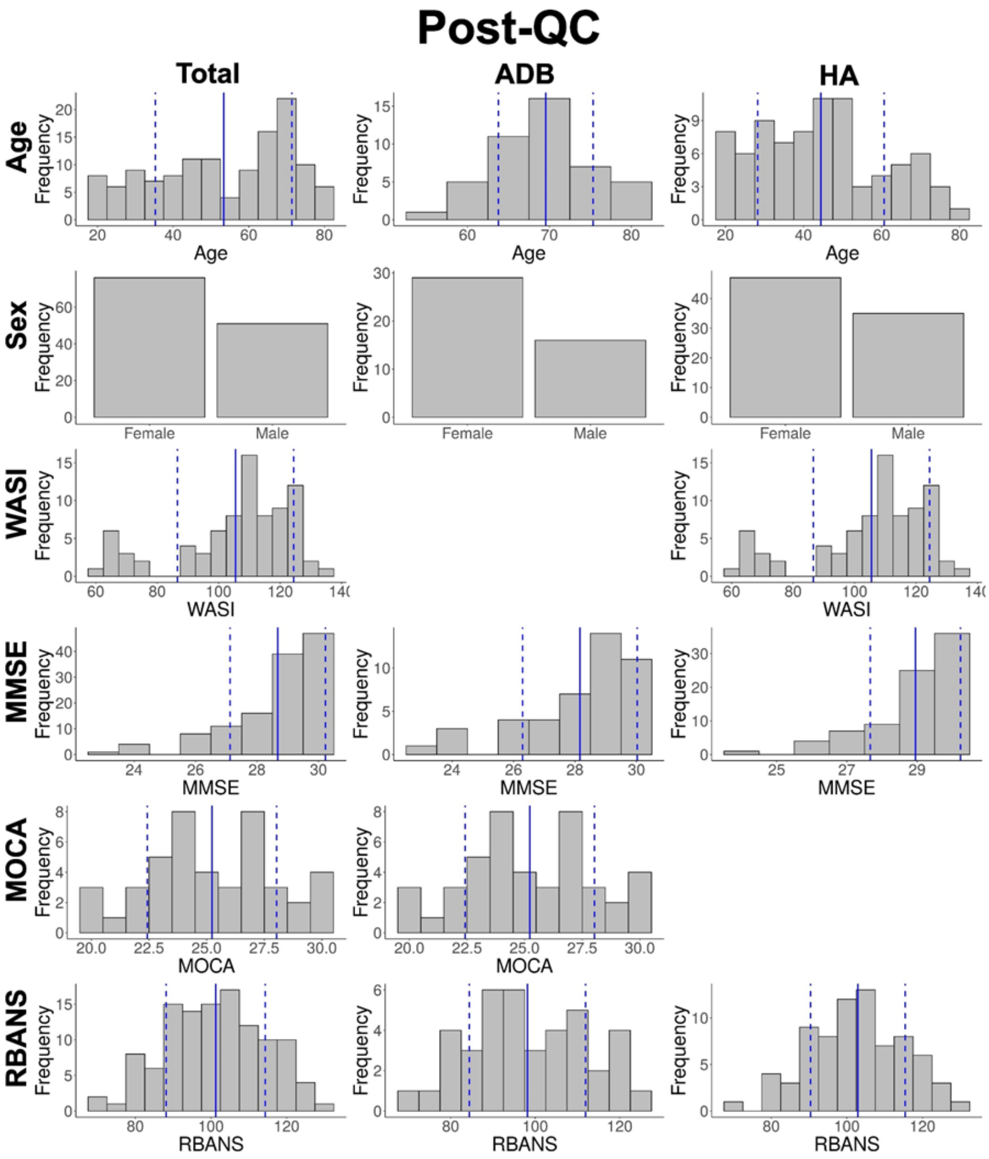
Post-QC
Total
ADB
HA
Age
Sex
WASI
MMSE
MOCA
RBANS
Frequency
Female
Male
Age
WASI
MMSE
MOCA
RBANS

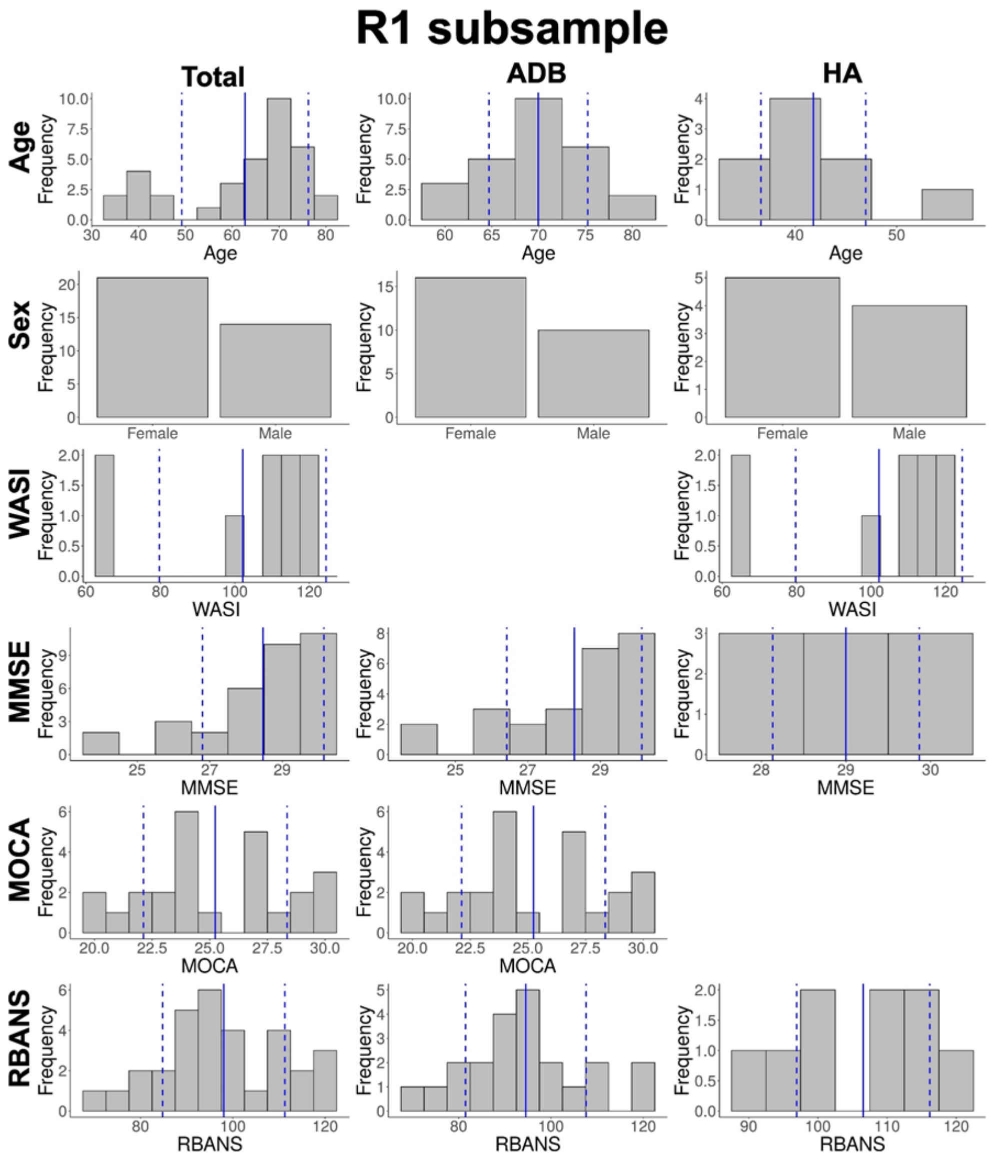


**Supplementary figure 1. Histograms of demographic variables.** Histograms for all variables in Table 1. The full blue line is the variable mean, with the dotted blue lines representing +/- 1 SD of the variable.

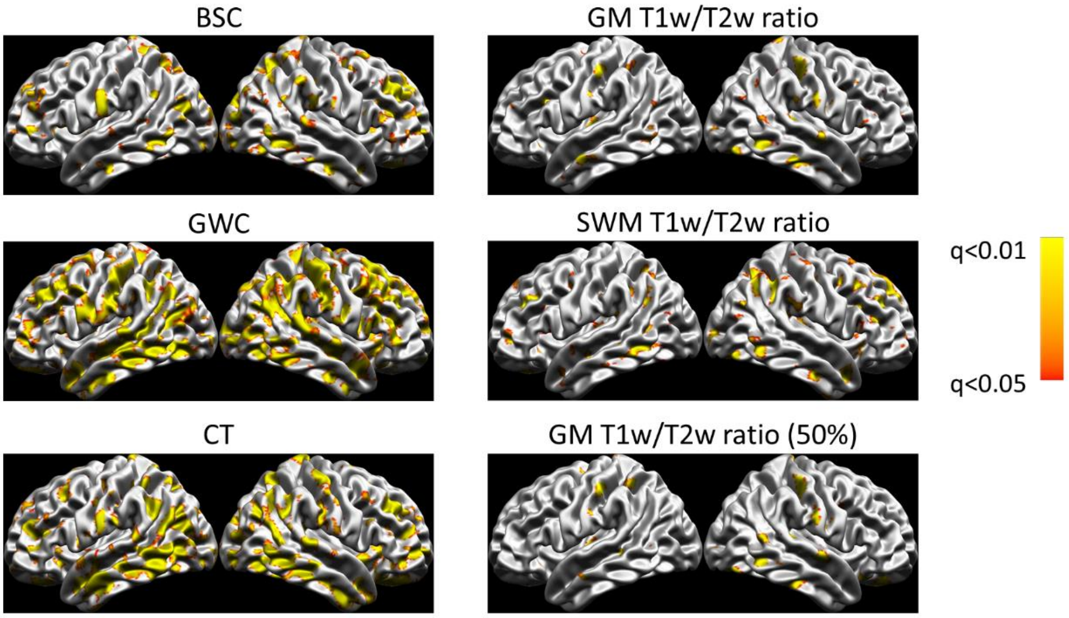


**Supplementary figure 2. Vertex-wise correlations between markers and mean curvature.** Correlations are thresholded at the FDR 0.05 level. The color is indicative of the q-values (FDR-corrected p-values), where the lighter colors reflect lower q-values and warmer colors reflect higher q-values.

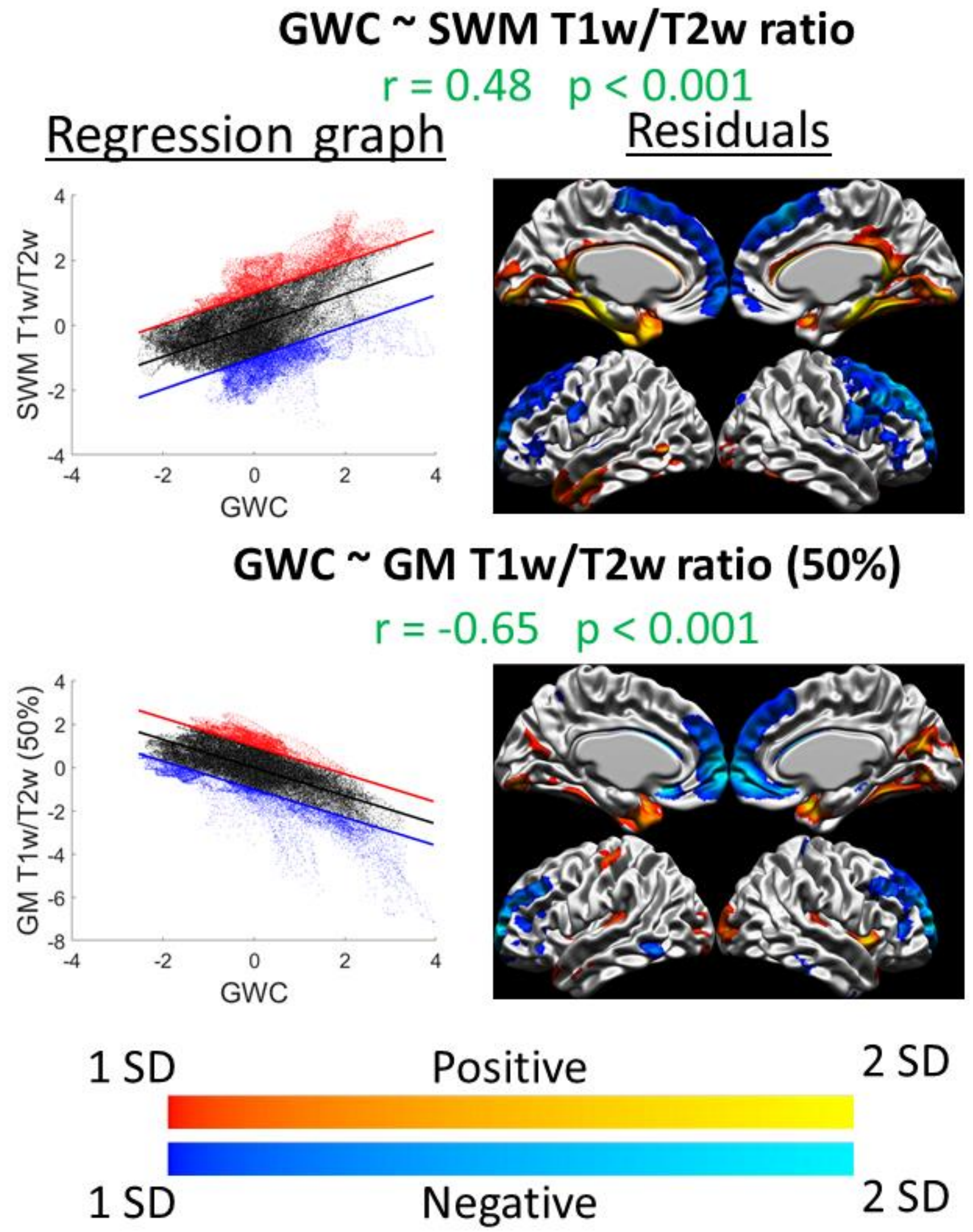


**Supplementary figure 3. Spatial distribution relationships of the linear age effect: graphs and residuals.** For each significant correlation, the left figure is the spatial regression in graph form, where the x-axis are the Z-scored values of the first marker, the y-axis are the Z-scored values of the second marker, the regression line is shown in black, and the +1 SD and -1 SD lines are shown in red and blue respectively (representing the thresholds set for values that are far from the regression line and exhibit less the observed relationship). The right figure is the vertex-wise residuals from the regression thresholded at +/- 1 SD (cold colors indicate vertices below the regression line in blue in the left graph and warm colors indicate vertices above the regression line in red in the left graph, and lighter colors indicate higher residual values and darker colors indicate lower residual values).

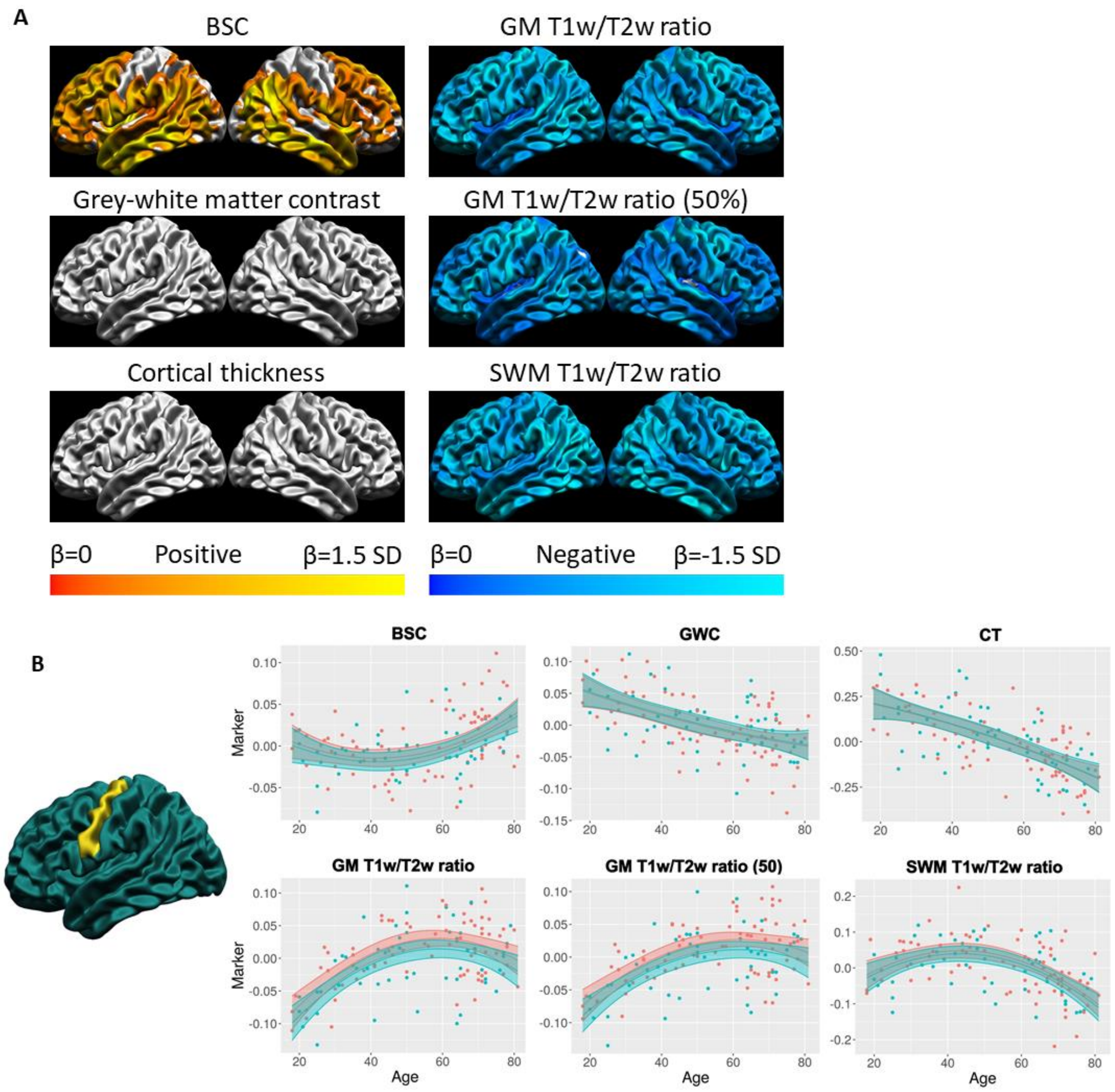


**Supplementary figure 4. Quadratic age trajectories. A.** For each marker, the mean and standard deviation of the age betas were calculated and used to threshold the colors. Cortical maps are thresholded for significance at the FDR 0.05 level. Cold colors indicate negative age betas and warm colors indicate positive age betas. Light colors indicate higher age betas relative to the marker mean, and dark colors indicate lower age betas. **B.** Example of the age trajectory of each marker at one vertex in the precentral gyrus. Blue observations represent male participants and red observations represent female participants. The x-axis is age and the y-axis is the marker value residualized for mean curvature.

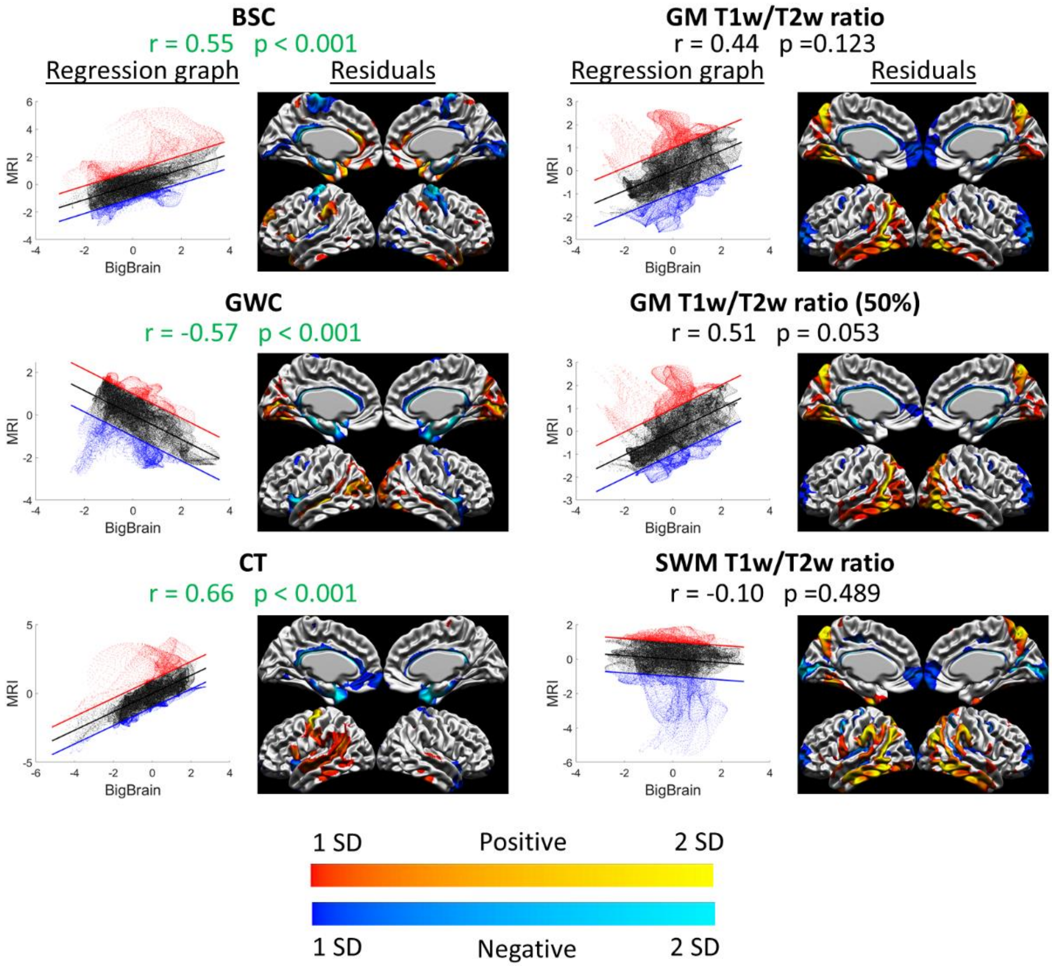


**Supplementary figure 5. MRI-BigBrain relationships: graphs and residuals.** For each significant correlation, the left figure is the spatial regression in graph form, where the x-axis are the Z-scored values of the first marker, the y-axis are the Z-scored values of the second marker, the regression line is shown in black, and the +1 SD and -1 SD lines are shown in red and blue respectively (representing the thresholds set for values that are far from the regression line and exhibit less the observed relationship). The right figure is the vertex-wise residuals from the regression thresholded at +/- 1 SD (cold colors indicate vertices below the regression line in blue in the left graph and warm colors indicate vertices above the regression line in red in the left graph, and lighter colors indicate higher residual values and darker colors indicate lower residual values).

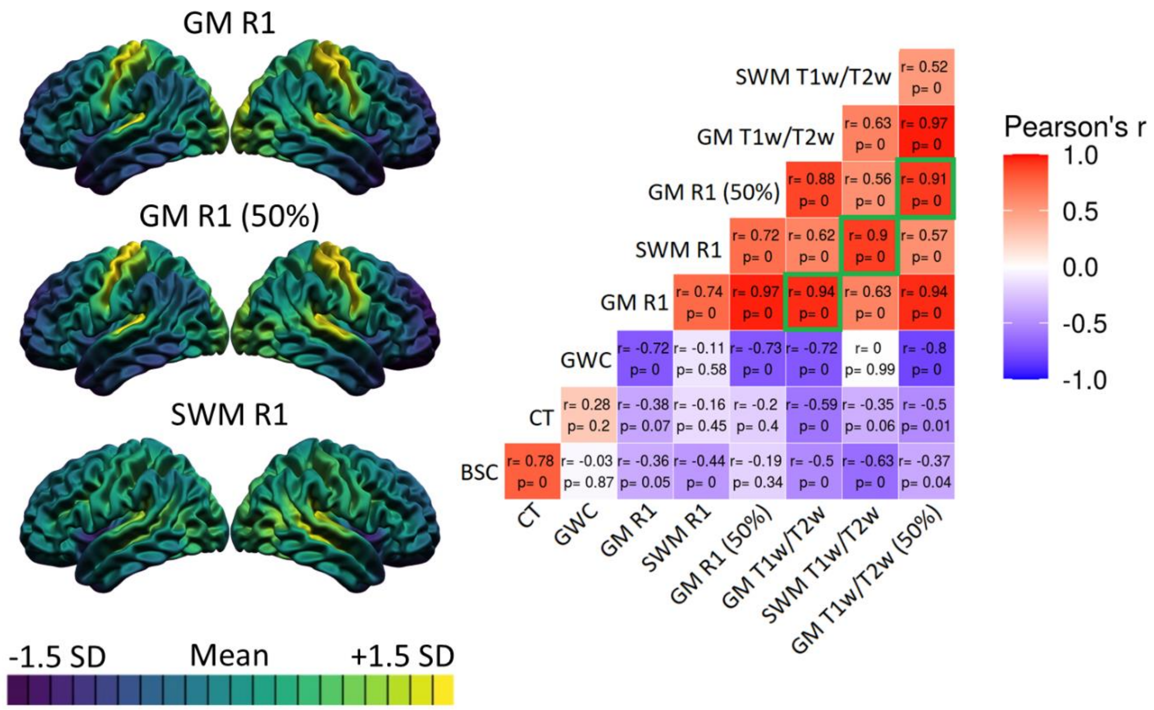


**Supplementary figure 6. Spatial distribution correlations including R1 measures.** For each R1 spatial distribution map, the mean and standard deviation of the surface were calculated and used to threshold the colors. Purple areas indicate lower values relative to the mean of that marker, while yellow areas indicate higher values. The correlation matrix includes Pearson's correlation coefficients (r) and FDR-corrected p-values. The color of each correlation block is linked to the correlation coefficient: positive coefficients are red and negative coefficients are blue, and high coefficients are more saturated and low coefficients tend towards white. Correlations between R1 and T1w/T2w ratio sampled at the same depth are highlighted with a green outline.

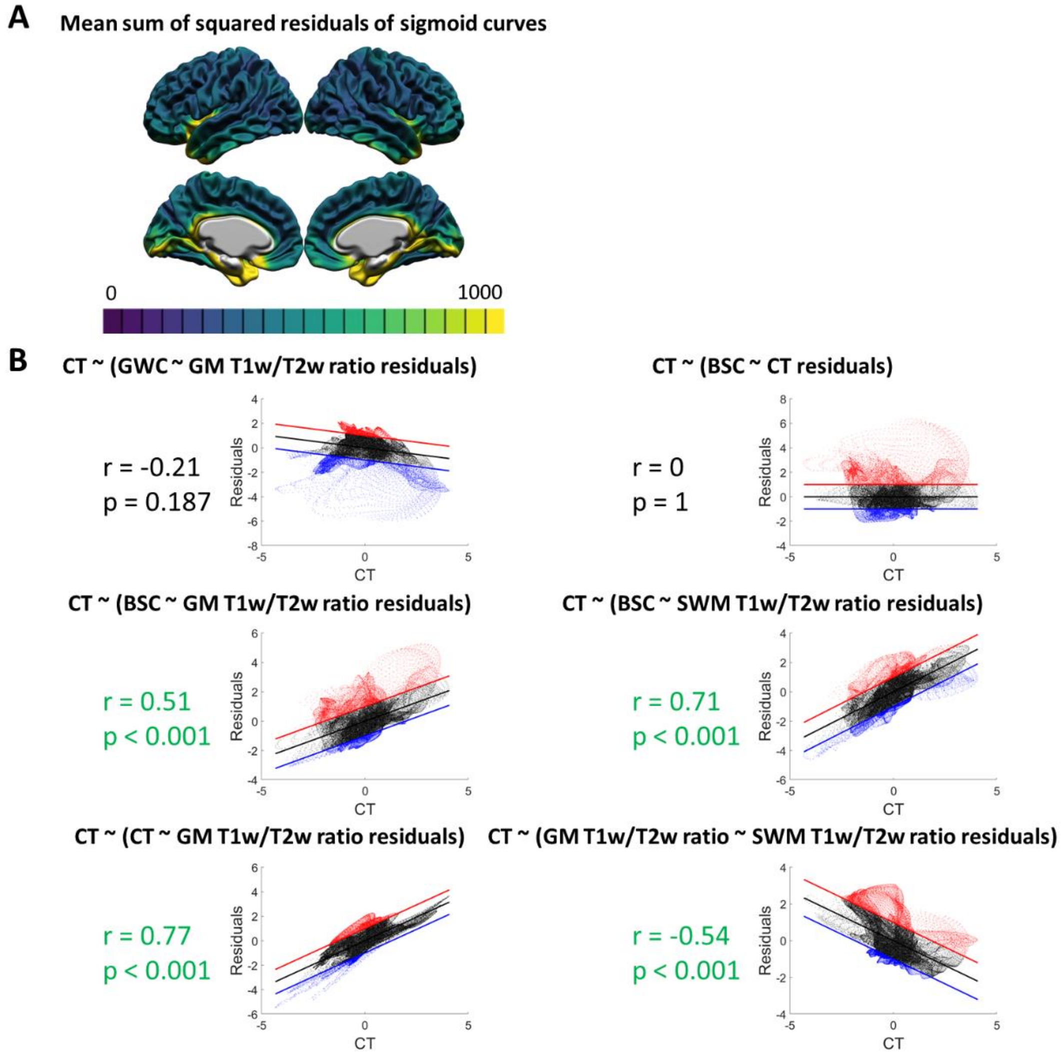


**Supplementary figure 7. Investigation of the cause of high residuals from spatial correlations in medial regions. A.** Vertex-wise mean sum of squared residuals of the sigmoid curves generated to compute the Boundary Sharpness Coefficient (BSC) marker. Lower values in purple indicate a better fit of the sigmoid curve to the cortical profile, while higher values in yellow indicate a worst fit of the sigmoid curve to the cortical profile. **B.** Correlations between cortical thickness (CT) and residuals from significant correlations between spatial distributions. Pearson's correlation coefficient (r) and p-values are highlighted in green if the correlation is significant at the 0.05 level. The right figure is the vertex plot of the correlation where the x-axis are the values of the first marker, the y-axis are values of the second marker and the regression line is shown in red.

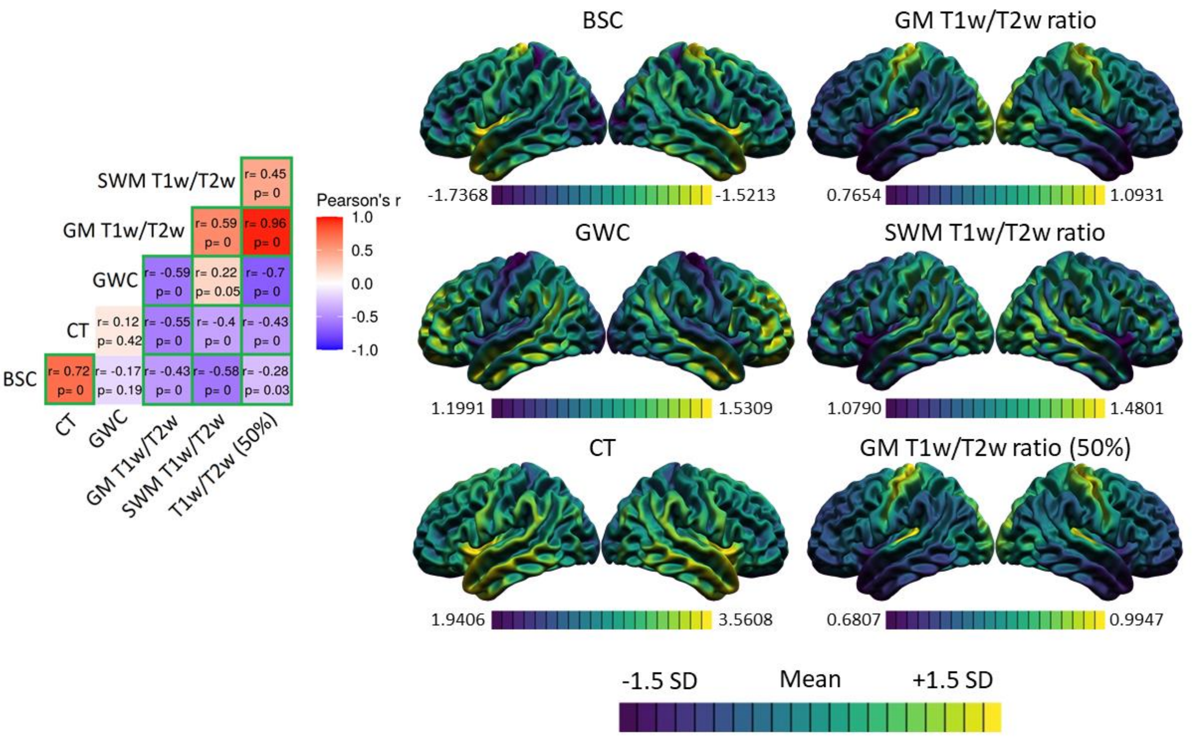


**Supplementary figure 8. Spatial distributions of the markers and correlations (5mm smoothing).** Same analysis as Figure 2, but with markers smoothed with a 5mm full-width half-max heat kernel. For each marker, the mean and standard deviation of the surface were calculated and used to threshold the colors. Purple areas indicate lower values relative to the mean of that marker, while yellow areas indicate higher values. The correlation matrix includes Pearson's correlation coefficients (r) and FDR-corrected p-values. The color of each correlation block is linked to the correlation coefficient: positive coefficients are red and negative coefficients are blue, and high coefficients are more saturated and low coefficients tend towards white. Significant correlations at the FDR 0.05 level are highlighted with a green outline.

**A**

| | BSC | GWC | CT | GM T1w/T2w | GM T1w/T2w (50%) | SWM T1w/T2w |
|---|---|---|---|---|---|---|
| **Linear** | 49% | 67% | 68% | 4% | 10% | 3% |
| **Quadratic** | 37% | 16% | 18% | 77% | 69% | 83% |
| **Cubic** | 14% | 17% | 14% | 19% | 20% | 14% |

**B**

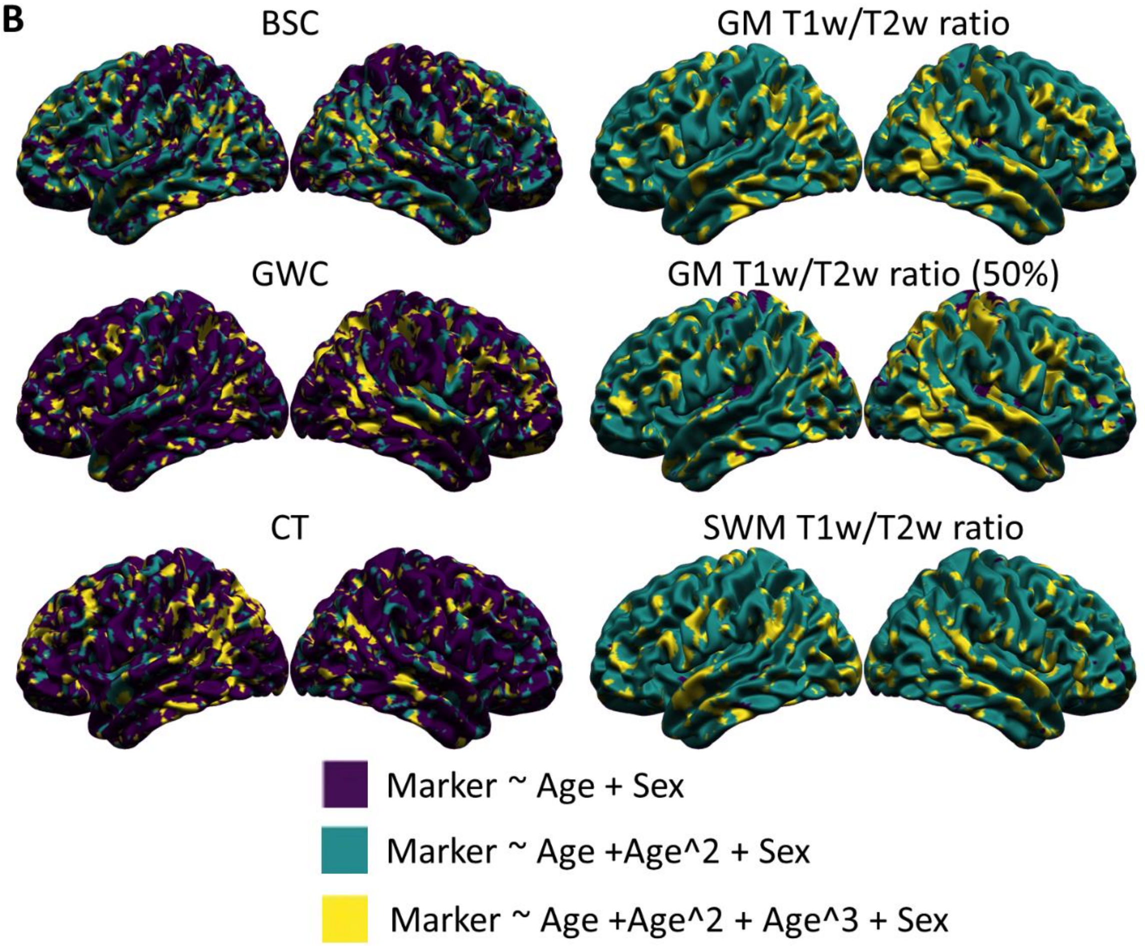


**Supplementary figure 9. Vertex-wise best age trajectory shape between linear, quadratic and cubic for each marker (5mm smoothing).** Same analysis as Figure 4, but with markers smoothed with a 5mm full-width half-max heat kernel applied after regressing out curvature. **A.** Table illustrating the proportion of vertices best fitted by each age model for each marker according to the Akaike Information Criterion (AIC), with the age model best fitting the highest proportion of vertices highlighted in green. **B.** Spatial distribution of the AIC results. Purple areas indicate a better fit of the linear age trajectory, green areas indicate a better fit of the quadratic age trajectory, and yellow areas indicate a better fit of the cubic age trajectory.

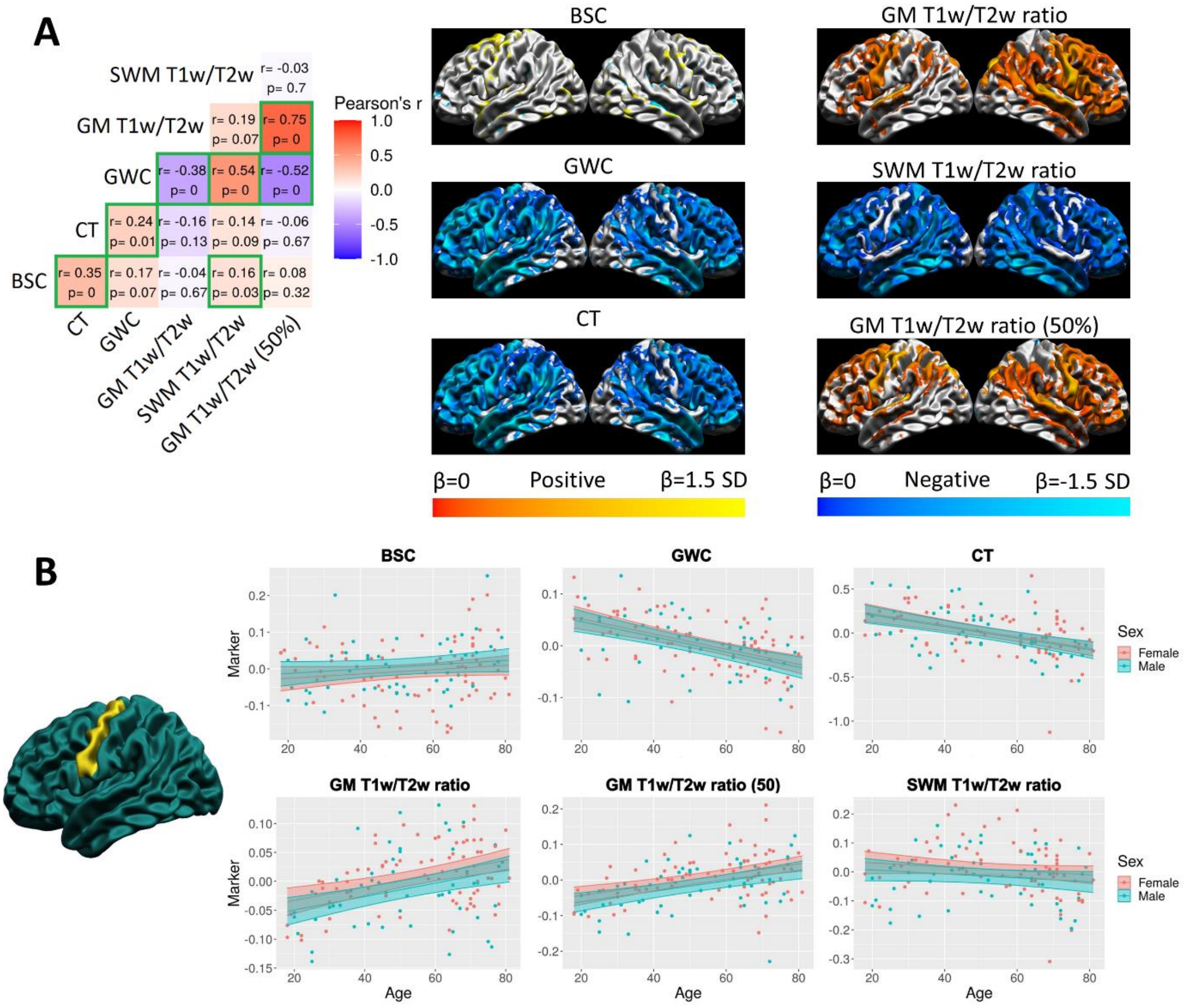


**Supplementary figure 10. Spatial distribution of the linear age effect of the markers and correlations (5mm smoothing).** Same analysis as Figure 5, but with markers smoothed with a 5mm full-width half-max heat kernel applied after regressing out curvature. **A.** For each marker, the mean and standard deviation of the age betas were calculated and used to threshold the colors. Cortical maps are thresholded for significance at the FDR 0.05 level. Cold colors indicate negative age betas and warm colors indicate positive age betas. Light colors indicate higher age betas relative to the marker mean, and dark colors indicate lower age betas. The correlation matrix includes Pearson's correlation coefficient (r) and FDR-corrected p-values. The color of each correlation block is linked to the correlation coefficient: positive coefficients are red and negative coefficients are blue, and high coefficients are more saturated and low coefficients tend towards white. Significant correlations at the FDR 0.05 level are circled in green. **B.** Example of the age trajectory of each marker at one vertex in the precentral gyrus where the age beta of each marker was significant at the FDR 0.05 level. Blue observations represent male participants and red observations represent female participants. The x-axis is age and the y-axis is the marker value residualized for mean curvature.

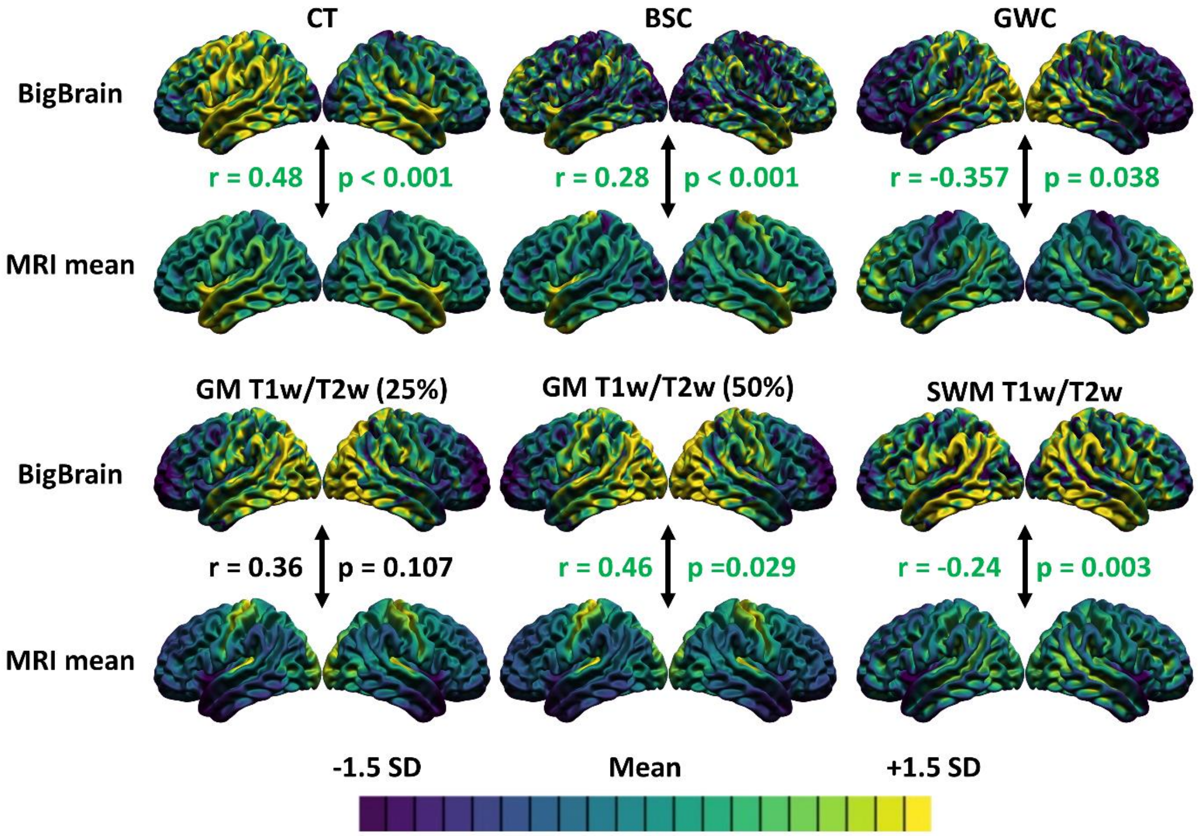


**Supplementary figure 11. Correlations between the spatial distributions of the markers generated on MRI and markers generated on BigBrain (5mm smoothing).** Same analysis as Figure 6, but with markers smoothed with a 5mm full-width half-max heat kernel. For each marker, the mean and standard deviation were calculated and used to threshold the colors. More specifically, purple areas indicate lower values relative to the mean of that marker, while yellow areas indicate higher values. The Pearson's correlation coefficient and p-value are colored green if the relationship is significant at the 0.05 level.

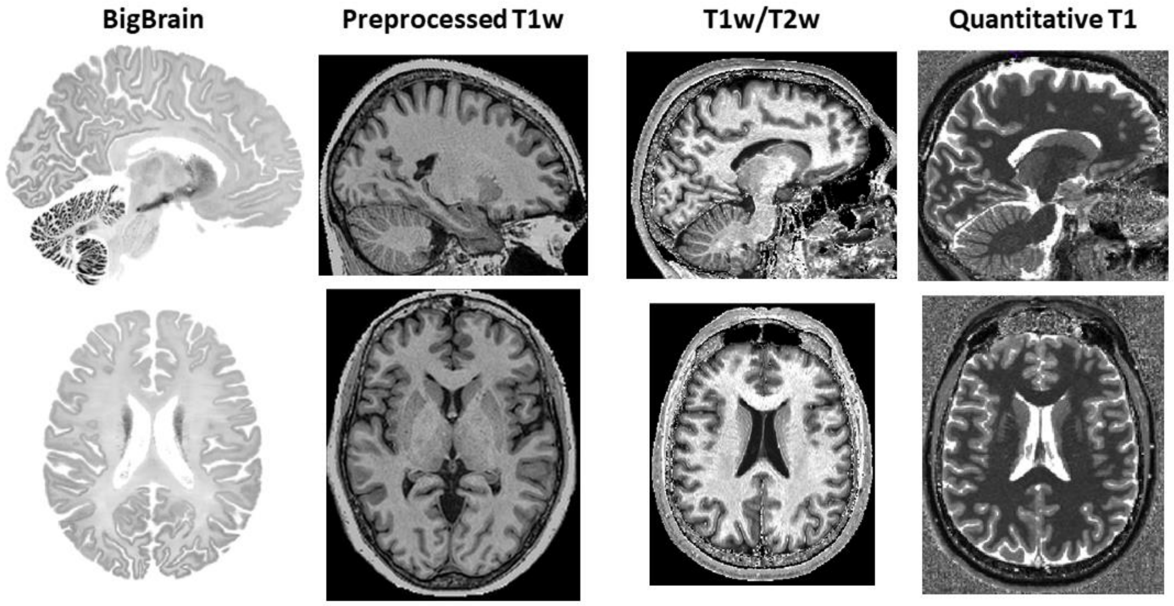


**Supplementary figure 12. Comparison of intensity between the BigBrain and MRI scans.** Sagittal (top row) and horizontal (bottom row) are located roughly in the middle of the respective brains. All MRI scans (preprocessed T1w, T1w/T2w ratio and quantitative T1) are from a 65 year old cognitively normal female.
